# Discovery of a Selective Inhibitor of ZIP14 with Therapeutic Potential for Cancer-associated Cachexia

**DOI:** 10.1101/2025.10.23.682519

**Authors:** Takafumi Hara, Gen Tanaka, Tomonori Tamura, Masaomi Terajima, Kengo Hamamura, Yuya Yoshida, Toru Kimura, Yusuke Kasai, Yuta Nakayama, Takumi Umeyama, Kohei Hosoi, Ayaka Noguchi, Yasuno Nakai, Atsushi Hijikata, Koji Matsukawa, Satoru Ujihara, Tetsuhiro Kawabe, Hiroki Taguchi, Hitomi Fujishiro, Supak Jenkitkasemwong, Kazuto Nunomura, Bangzhong Lin, Ayako Fukunaka, Emi Yoshigai, Kenji Mishima, Shinsaku Nakagawa, Michell D. Knutson, Hiroshi Imagawa, Naoya Matsunaga, Shigehiro Ohdo, Itaru Hamachi, Hiroyuki Sakurai, Toshiyuki Fukada

## Abstract

ZIP14/SLC39A14, a membrane-bound metal transporter, is essential for systemic metal homeostasis and has been implicated in inflammatory and metabolic disorders, including cancer-associated cachexia. Despite its biological and therapeutic significance, no selective inhibitors have been identified. Here, we identify 1-phenyl-8-(2-phenylethyl)-1,3,8-triazaspiro[4.5]decan-4-one (PPTD) as the first selective small-molecule inhibitor of ZIP14. PPTD efficiently blocks ZIP14-mediated uptake of zinc, iron, manganese, and cadmium, while sparing the closely related transporter ZIP8/SLC39A8. Mechanistically, PPTD binds specifically to a pocket formed at the dimer interface of ZIP14, as revealed by AlphaFold3 structural prediction, ligand–interaction profiling, structure–activity analyses, and site-directed mutagenesis, providing direct evidence for a targeted inhibition mechanism. ZIP14-driven metal influx promotes reactive oxygen species and lipid peroxidation, leading to cytotoxicity, which PPTD effectively reverses. *In vivo*, PPTD ameliorates major features of cancer cachexia in mice, including weight loss, reduced survival, muscle wasting, impaired locomotor activity, and disease progression. PPTD thus provides both a chemical probe to dissect ZIP14 function and a potential therapeutic candidate for cancer cachexia, establishing a foundation for the development of therapies targeting ZIP14-mediated metal dysregulation.

## Introduction

Essential trace elements are critical in maintaining vital physiological functions and are indispensable for various life processes ^1^. Of these elements, iron is the most abundant in humans, followed by zinc. They are widely distributed biological metals, integral to a vast array of enzymes and metalloproteins, and involved in numerous biochemical pathways. Therefore, dysregulation in the quantity of these metals impairs physiological functions. The regulation of intracellular zinc concentrations is controlled by two primary families of metal transporters encoded by the SLC30A1–10 and SLC39A1–14, respectively: ZnTs (zinc transporters) and ZIPs (Zrt/Irt-like proteins or solute carrier 39 or SLC39) ^2^. They function as homo- or heterodimers to coordinate the maintenance of intracellular zinc homeostasis ^3–5^. ZnTs function primarily as efflux transporters, exporting excess zinc from the cytoplasm, whereas ZIPs mediate the influx of zinc into the cytosol, ensuring homeostatic zinc concentrations ^6^.

Recent advances in genetic research have uncovered strong associations between ZIP transporters and various human diseases ^7^. For example, loss-of-function mutations in ZIP4 cause acrodermatitis enteropathica (AE), an autosomal recessive disorder characterized by impaired zinc absorption, growth retardation, immune dysfunction, and dermatitis ^8^. Hypomorphic mutations in ZIP7 in patients with agammaglobulinemia and early-onset infections underscore its essential role in B cell development and acquired immunity ^9^. ZIP13 is critical for connective tissue homeostasis, including bone, skin, and teeth. Loss-of-function mutations in ZIP13 lead to Ehlers-Danlos syndrome, spondylodysplastic type 3 (EDSSPD3), an autosomal recessive connective tissue disorder ^10,11^. Systemic ZIP14 deficiency in mice results in growth retardation and skeletal abnormalities ^12^. In humans, loss-of-function mutations in ZIP14 have been linked to neurodegenerative disorders such as parkinsonism ^13^, most likely due to impaired manganese regulation ^14^, highlighting the critical role of ZIP14 in maintaining trace metal homeostasis and physiological integrity.

ZIP14 was initially identified as a plasma membrane-associated zinc transporter ^15^. Subsequently, it was found to facilitate the transport of other divalent metals, including manganese, cadmium ^16,17^, and iron ^18,19^. Iron is the most abundant essential trace element; however, to avoid iron overload, cells suppress transferrin receptor expression once intracellular iron levels reach sufficient concentrations. Under such conditions, non-transferrin-bound iron (NTBI), a redox-active and potentially toxic form, accumulates in the extracellular milieu and is taken up by cells via alternative transporters, including DMT1 and ZIP14 ^20^. Consequently, the activity of ZIP14 has been implicated in metal-related pathologies. For instance, in hereditary hemochromatosis, iron overload leads to elevated levels of NTBI. ZIP14 facilitates NTBI uptake into hepatocytes, aiding in iron clearance from circulation. *Zip14*-deficient mice are resistant to hepatic and pancreatic iron overload, substantiating its central role in systemic iron regulation (21). More recently, gain-of-function of ZIP14, through elevated expression, has been linked to cancer-associated cachexia, a complex syndrome characterized by progressive skeletal muscle wasting. Inflammatory cytokines upregulate ZIP14 in cachectic muscle, resulting in excessive zinc influx into muscle progenitor cells, impaired myogenesis, and disease progression ^22^. These findings highlight the critical role of ZIP14 regulation in maintaining metal homeostasis and overall physiological balance.

The pivotal roles of metal transporters in several pathological processes have positioned them as attractive targets for pharmacological intervention ^2^. A low molecular weight inhibitor targeting ZIP7, namely NVS-ZP7-4, was identified by a screen intended to identify Notch signaling inhibitors. Inhibition of ZIP7 results in the accumulation of zinc within the endoplasmic reticulum (ER), thereby inducing ER stress, disrupting Notch signaling, and triggering apoptosis ^23^. Another approach has involved fragment-based drug discovery aimed at DMT1. Screening of the GDB-17 fragment library led to the identification of trifluoromethylsulfone and thiophene carboxylic acid as weak inhibitors of DMT1; tetrahydrocarbazole was identified as a nonspecific ZIP8 inhibitor ^24^. To date, however, these compounds have not progressed to clinical or pharmacological application stages.

Given ZIP14’s involvement in various pathophysiological processes, it represents a promising therapeutic target for diseases characterized by dysregulated metal homeostasis. Here, we report the discovery of 1-phenyl-8-(2-phenylethyl)-1,3,8-triazaspiro[4.5]decan-4-one (PPTD), the first identified selective small-molecule inhibitor of ZIP14 with therapeutic potential.

## Results

### Identification of a specific small-molecule inhibitor of ZIP14

We performed a high-throughput screen to identify small-molecule inhibitors of ZIP14-mediated zinc uptake (Fig. 1A), using TREx-human ZIP14 (hZIP14) and ZIP8 (hZIP8) cells where human ZIP14 or ZIP8 was induced by tetracycline (Tet) treatment, respectively (Supplementary Fig. 1). After three systematic rounds of screening, including the elimination of false positives such as zinc chelators, fluorescent quenchers, and esterase inhibitors, PPTD (Fig. 1B) was identified as a potent ZIP14 inhibitor. PPTD exhibited an IC₅₀ of 8.1 μM for ZIP14-mediated zinc uptake in TREx-hZIP14 cells, whereas no significant inhibition was observed for ZIP8, the closest paralog of ZIP14, up to 37μM in TREx-hZIP8 cells (Supplementary Fig. 2), implicating PPTD as a specific inhibitor of ZIP14.

**Fig. 1.**
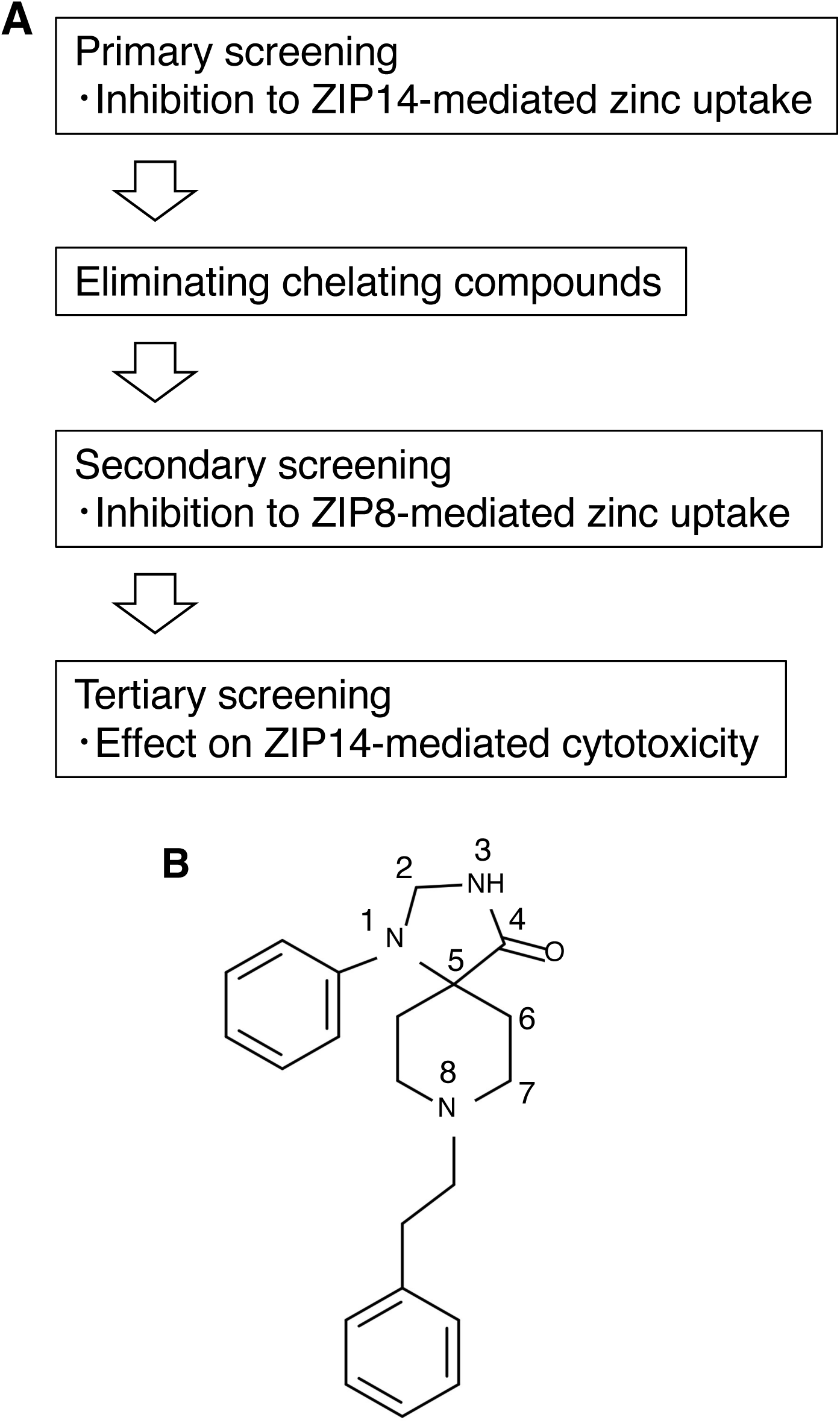
High-throughput screening for ZIP14 inhibitors. **A** Schematic workflow of the high-throughput screening to identify ZIP14 inhibitors. **B** Chemical structure of 1-phenyl-8-(2-phenylethyl)-1,3,8-triazaspiro[4.5]decan-4-one (PPTD).

### PPTD selectively inhibits ZIP14-mediated metal uptake and cytotoxicity

ZIP14 facilitates the transport of multiple divalent metals, including zinc, iron, manganese, and cadmium ^17^. We conducted radiotracer uptake assays in both TREx-cell lines (Supplementary Fig. 1) and *Xenopus laevis* oocytes (Supplementary Fig. 3) expressing human ZIP14 to evaluate the effect of PPTD on ZIP14-mediated metal transport. PPTD significantly inhibited the uptake of all four metals (Fig. 2A–D). PPTD exhibited an equivalent IC₅₀ against both ZIP14-mediated zinc and iron uptake assessed by *Xenopus laevis* oocytes system (Supplementary Table 1), suggesting its inhibition of metal transport by the same or similar mechanism. We examined PPTD’s effect on ZIP8, a closely related metal transporter, to assess its selectivity. In both *Xenopus* oocytes expressing ZIP8 and TREx-hZIP8 cells, PPTD did not affect the transport of zinc, iron, manganese, or cadmium (Fig. 2E–H), confirming that PPTD could selectively target ZIP14.

**Fig. 2.**
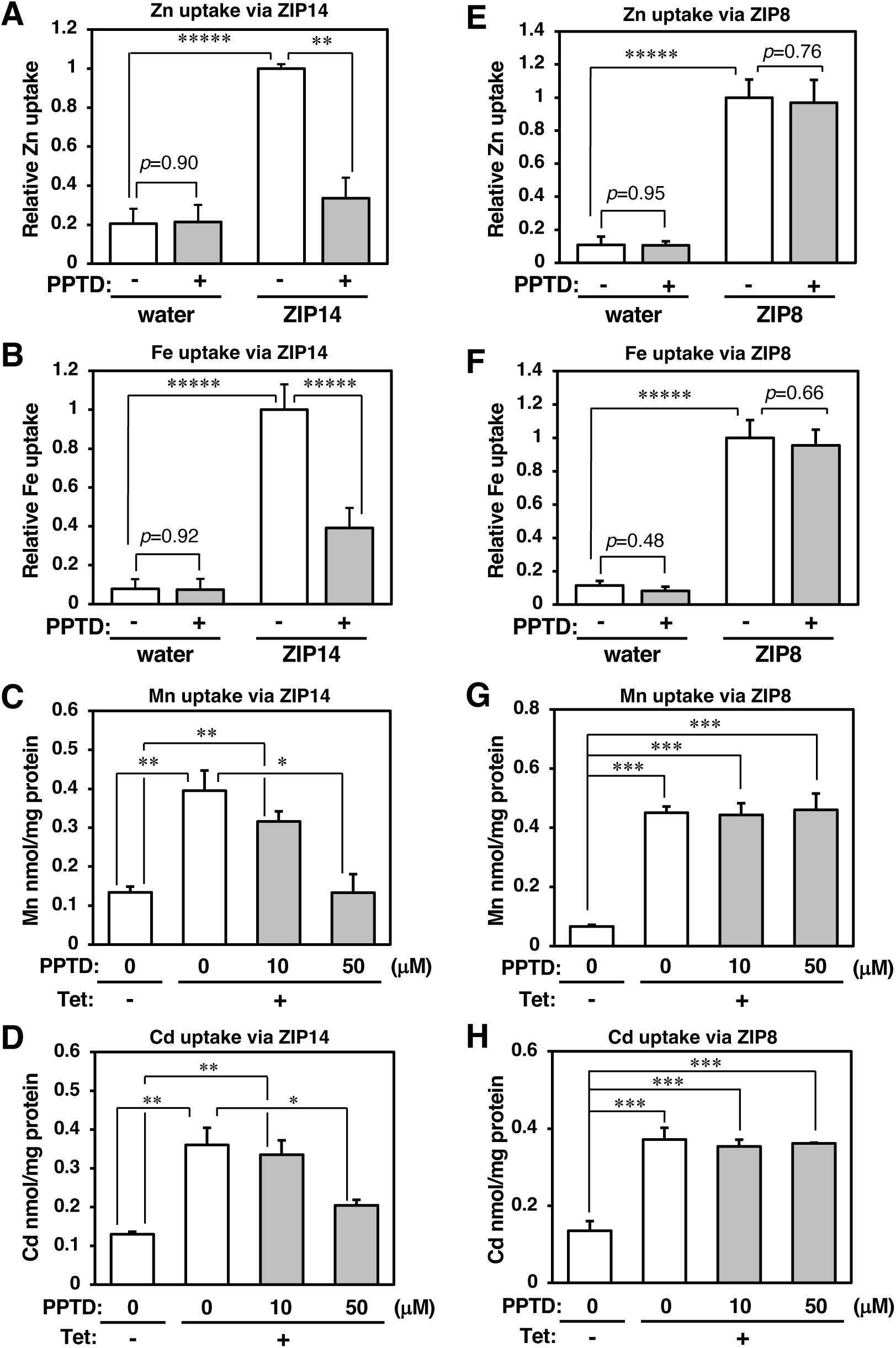
PPTD selectively inhibits ZIP14-mediated metal transport. *Xenopus* oocytes injected either with (**A, B**) human ZIP14 or (**E, F**) human ZIP8 cRNA were incubated with either (**A, E**) ⁶⁵Zn or (**B, F**) ⁵⁵Fe in the presence of (**A, E**) 50 µM or (**B, F**) 10 µM PPTD for 30 min at 22 °C. Radioactivity was measured as described under *Methods*. Data are normalized against DMSO-treated transporter-expressing controls, and presented as mean ± standard deviation (S.D.) (n=3 independent experiments). Similarly, (C, D) TREx-hZIP14 or (G, H) TREx-hZIP8 cells were treated with Tet (1 μg/mL) for 24 h, followed by incubation either with (**C, G**) ⁵⁴MnCl₂ or (D, H) ¹⁰⁹CdCl₂ in the presence of PPTD at the indicated concentrations for 1 h. Radioactivity was measured as described under *Methods*. The data (mean + SD) represent results from three independent experiments. Statistical significance was determined using Student’s *t*-test (A, B, E, F) or one-way ANOVA followed by Dunnett’s post hoc test (C, D, G, H). (\**p* < 0.05, \*\**p* < 0.01, \*\*\**p* < 0.005, \*\*\*\**p* < 0.001, \*\*\*\*\**p* < 0.0005).

Given that metal overload induces cytotoxicity ^25,26^, we assessed whether PPTD could mitigate ZIP14-mediated metal toxicity. Induced ZIP14 expression in TREx-hZIP14 cells sensitized them to zinc, iron, manganese, and cadmium, all significantly attenuated by PPTD treatment (Fig. 3A–D,). In contrast, no such protective effect was observed in TREx-hZIP8 cells (Fig. 3E–H), reinforcing PPTD’s ZIP14-specific action. Further, ZIP14-dependent metal uptake elevated reactive oxygen species (ROS) and lipid peroxides (LPO) generation, leading to cytotoxicity, which were significantly reduced by PPTD treatment (Fig. 4A and B). Similar results were obtained in mouse myoblast cells, C2C12 (Supplementary Fig. 4), supporting the effective blocking of ZIP14 functions by PPTD. These findings demonstrated that PPTD blocked ZIP14-mediated metal influx and attenuated its associated cytotoxicity, underscoring its potential as a selective pharmacological tool.

**Fig. 3.**
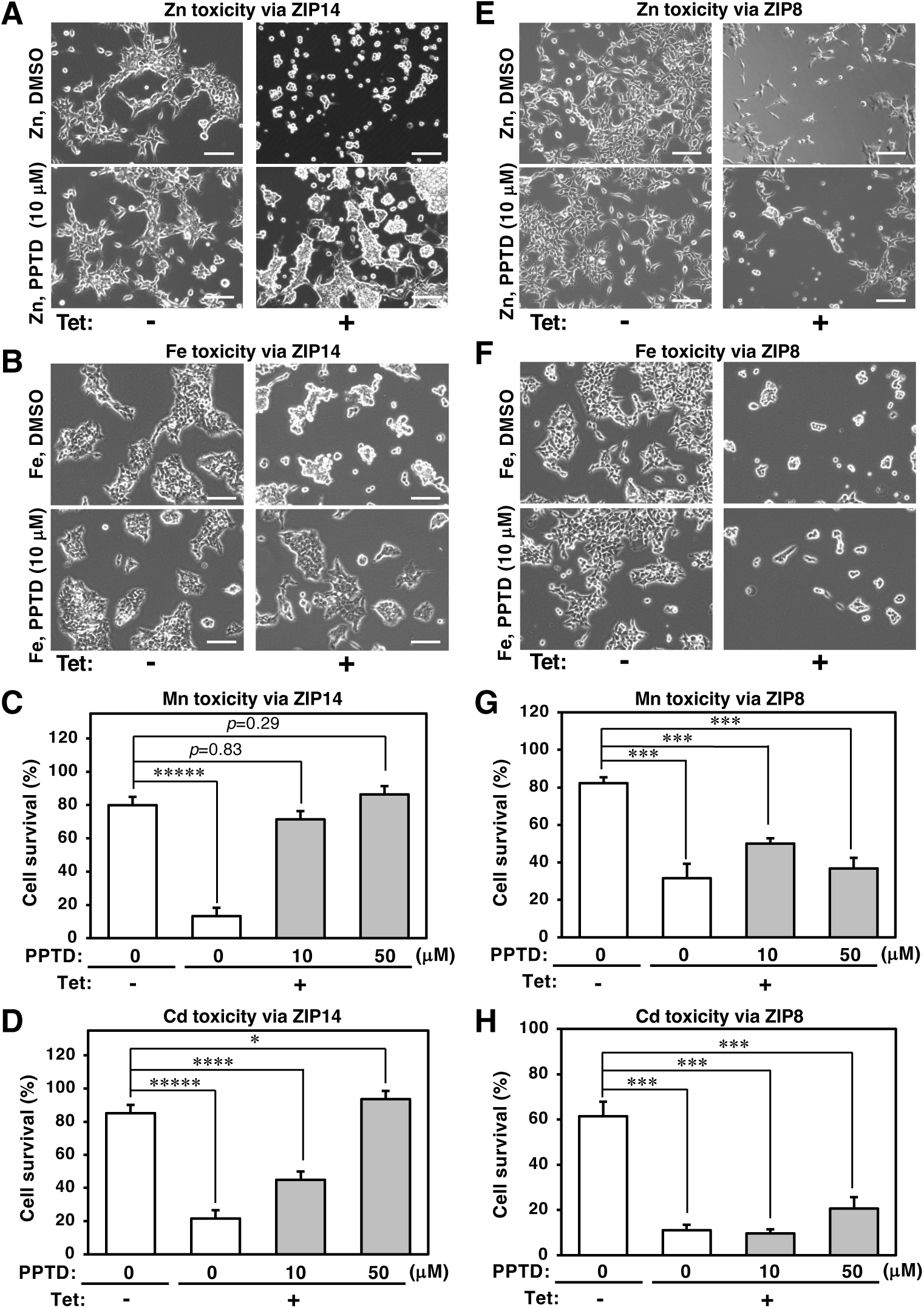
PPTD inhibits ZIP14-mediated cytotoxicity. TREx-hZIP14 cells were treated with Tet (1 μg/mL) in combination with (**A**) ZnSO_4_ (100 μM), (**B**) FeSO_4_ (20 μM), (**C**) MnCl₂ (100 μM), or (**D**) CdCl₂ (20 μM), with or without PPTD at the indicated concentrations. Similarly, TREx-hZIP8 cells were treated under the same conditions (**E–H**). Phase-contrast images showing cell morphology (**A, B, E, F**). Cell viability (**C, D, G, H**) was quantified as described under *Methods*, and representative data (mean + SD) from three independent experiments are presented. Statistical significance was determined using one-way ANOVA followed by Dunnett’s post hoc test (**C, D, G, H**). (\**p* < 0.05, \*\*\**p* < 0.005, \*\*\*\**p* < 0.001, \*\*\*\*\**p* < 0.0005). Scale bar: 20 μm.

**Fig. 4.**
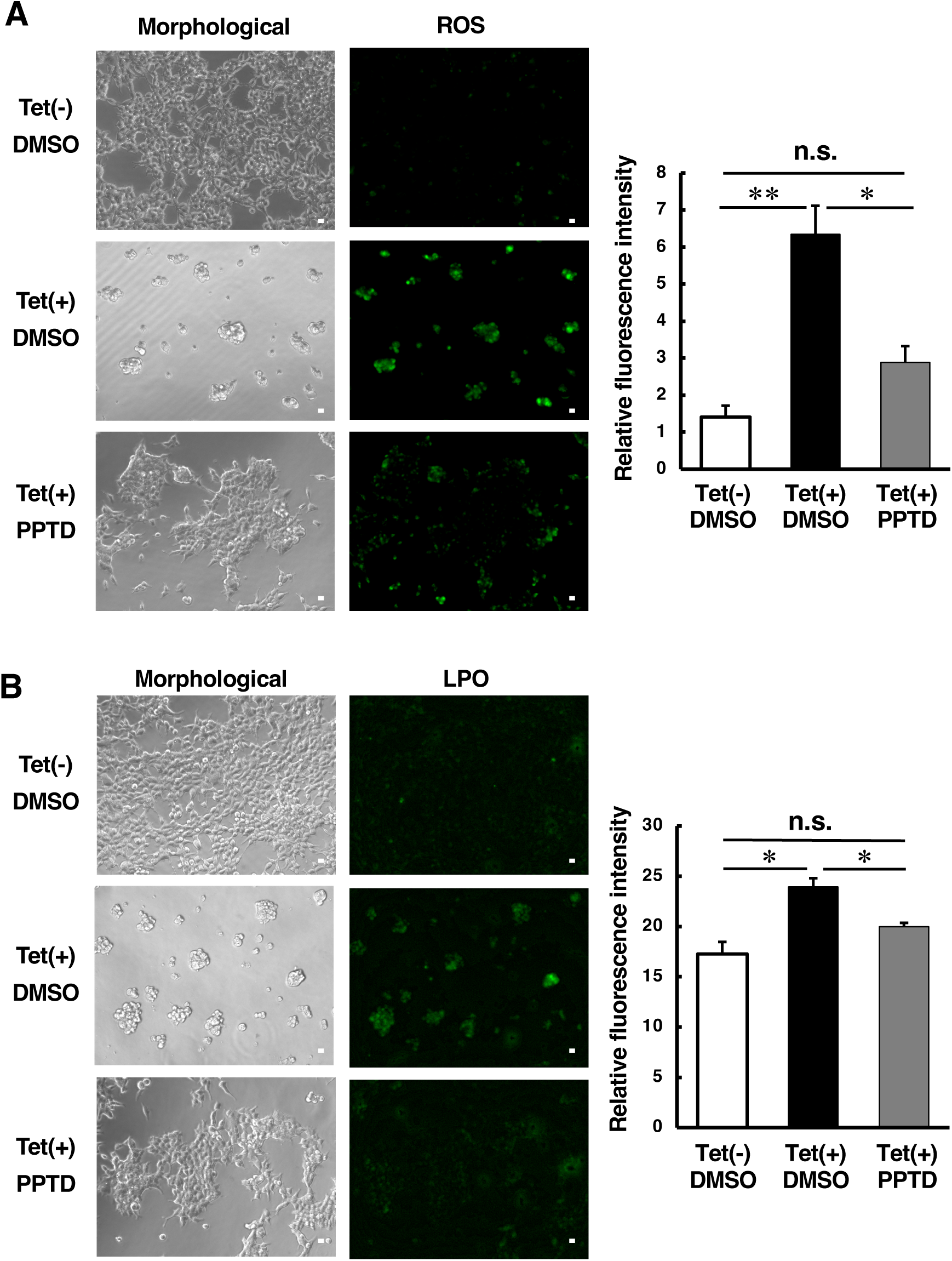
PPTD inhibits ZIP14-mediated generation of reactive oxygen species (ROS) and lipid peroxides (LPO) TREx-hZIP14 cells were treated with tetracycline (Tet, 1 μg/mL) for 48 h. They were then pretreated with PPTD, and both phase-contrast (morphological) and fluorescence images were captured to assess (**A**) ROS and (**B**) LPO levels (left panels), following the procedures described in the *Methods* section. Fluorescence intensity was subsequently quantified (right panels). Statistical significance was determined using a two-tailed unpaired Student’s *t*-test (\**p* < 0.05, \*\**p* < 0.01). Scale bar: 10 μm.

### SAR analysis for PPTD

We performed a structure–activity relationship (SAR) analysis to elucidate the structural determinants of PPTD activity, focusing on three major substitution sites: the 1-phenyl group at N1 (R1), the 2-phenylethyl group at N8 (R2), and the substituent at N3 (R3) (Fig. 1B).

At the R1 site (Supplementary Table 2), *p*-fluorination (N1-2) had little effect, whereas *p*-methoxylation (N1-1) or substitution with a methyl group (N1-3) abolished its activity, indicating the critical role of the unmodified 1-phenyl ring. At the R3 site (Supplementary Table 3), methylation (N3-1) preserved activity, but bulkier substituents (e.g., N3-2 and N3-3) eliminated inhibition, suggesting a steric constraint at this position. At R2 (Supplementary Table 4), both shortening (N8-2) or removing (N8-1) the 2-phenylethyl chain abrogated activity. The addition of large phenyl moieties (e.g., N8-12, N8-13, N8-17) disrupted the function. However, smaller modifications, including methyl, hydroxyl, sulfonamide, amine, azide, and fluoride groups, were tolerated. Notably, replacing the phenyl ring with a pyridine (N8-14, N8-15, N8-16) or a cyclohexyl group (N8-7) retained activity, suggesting limited flexibility in electronic and steric properties at this site.

These findings define a core pharmacophore: a conserved 1-phenyl ring at N1, a properly spaced 2-phenylethyl chain at N8, and minimal substitution at N3 are required for ZIP14 inhibition. Modifications to the 2-phenylethyl chain and the benzyl ring at N8 disrupted the inhibitory effect on ZIP14, suggesting that the structural features associated with these three major substitution sites, particularly N8, may be critical for PPTD associated with ZIP14 inhibition.

### Dimerization-dependent formation of a ZIP14-specific PPTD binding pocket

We modeled the ZIP14-PPTD complex using AlphaFold3 to clarify the molecular basis of ZIP14 inhibition by PPTD, predicting that PPTD is localized to the ZIP14 surface in the monomeric state (Fig. 5A left, and Supplementary Movie 1). However, upon dimerization of ZIP14, the compound was encapsulated within an internal pocket (Fig. 5A right, and Supplementary Movie 2). Structural overlay of the ZIP14-PPTD and ZIP14-zinc complexes could be predicted (Fig. 5B, and Supplementary Movie 3), suggesting that dimerization induced a conformational rearrangement creating defined ligand-binding pocket that obstructs the metal- binding and the transporting pathway (Supplementary Fig. 5A).

**Fig. 5.**
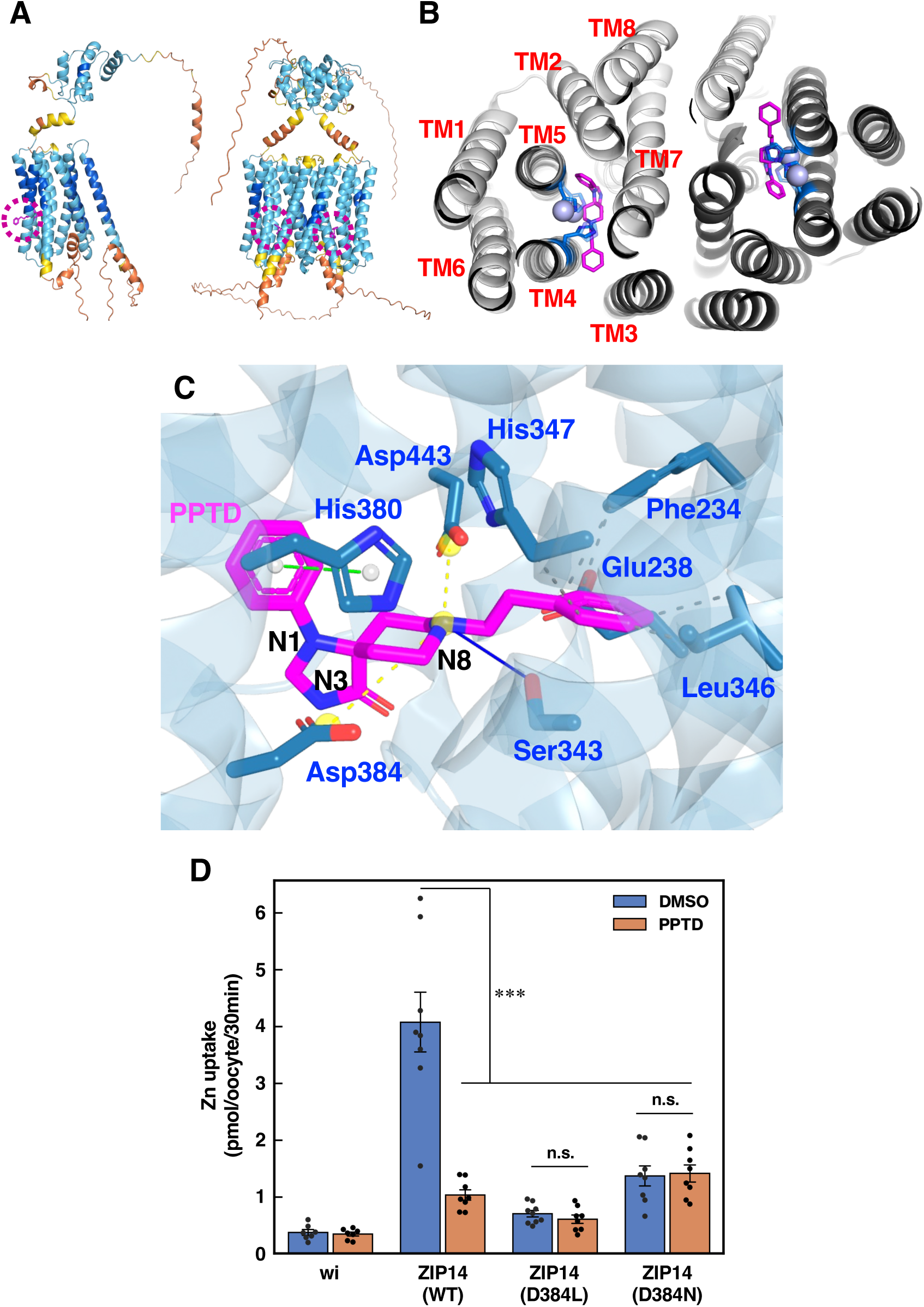
Identification of amino acids in ZIP14 involved in its association with PPTD. **A** Structural predictions of ZIP14-PPTD complexes using both monomeric (left) and dimeric (right) forms of human ZIP14 by AlphaFold3. Magenta circles indicate the predicted PPTD binding sites. Protein models are colored by structure prediction confidence estimated by predicted Local Distance Difference Test (pLDDT) (dark blue, pLDDT >90, light blue, pLDDT of 90-70, yellow, pLDDT of 70-50, orange, pLDDT <50). **B** Structural overlay of the ZIP14-PPTD and ZIP14-zinc complexes, predicted using AlphaFold3. PPTD and zinc bind in close proximity within the dimerized ZIP14 protein. magenta: PPTD, purple: zinc, blue: His347 and His380 residues. **C** Prediction of human ZIP14 amino acid residues interacting with PPTD by Protein–Ligand Interaction Profiler and AlphaFold3 (Supplementary Table 5). magenta: PPTD, green dot line: pi-stacking, yellow dot line: salt bridges, gray dot line: hydrophobic interactions, blue line: hydrogen bond. **D** Asp348 is essential for ZIP14-mediated zinc transport. *Xenopus* oocytes were injected with human ZIP14 (WT), ZIP14 (D348L), ZIP14 (D348N) cRNA, or water (wi, control). The uptake assay was performed with 10 μM ZnSO_4_ including a trace amount of radioactive tracer in the presence of 50 μM PPTD for 30 min at 22 °C. Radioactivity was measured as described under *Methods*. Statistical analysis was performed by one-way ANOVA followed by Tukey’s multiple comparison test. (\*\*\**p* < 0.001). All data are shown as mean ± S.E.M. (n= 6 to 9 oocytes).

We used the protein-ligand interaction profiler (PLIP) to characterize the binding interface (Supplementary Table 5), which identified several key residues interacting with PPTD (Fig. 5C and Supplementary Movie 4). Among them, His347 in transmembrane helix 4 (TM4) and His380 in TM5 contributed to PPTD binding through hydrophobic interactions and pi-stacking, respectively (Supplementary Table 5), which is consistent with the loss of inhibitory activity when the benzene ring is removed from the N1 and N8 positions of PPTD (Supplementary Tables 2 and 4). These amino acid residues are highly conserved among ZIP family members and coordinate zinc ions, forming a binuclear metal center (BMC) which is essential for structural stability and selective metal transport of ZIP members ^27^. A site-directed mutation in another PPTD-binding amino acid residue, Asp384 (Supplementary Table 5), located near the zinc-coordination HEXPHE motifs ^27^, disrupted zinc transport (Fig. 5D), suggesting the critical role of PPTD-binding amino acid residues within or near BMC for ZIP14-mediated zinc transport.

We also identified several key residues forming a high-affinity pocket unique to dimerized ZIP14 (Fig. 6A upper, Supplementary Fig. 5B, C, and Supplementary Movie 5). This pocket was structurally changed by the substitution of Ser343 with Tyr (S343Y) in ZIP14 (Fig. 6A lower and Supplementary Movie 6), which completely lost the transport activity (Fig. 6B). This pocket in dimerized ZIP14 was absent in dimerized ZIP8 (Supplementary Fig. 5D), potentially explaining the compound’s isoform specificity. We substituted Cys310 in ZIP8 with Gly (C310G) at the corresponding residue to evaluate whether this binding pocket could be engineered into ZIP8. Structural modeling of ZIP8 (C310G) mutant revealed a newly formed PPTD-binding pocket (Supplementary Fig. 6A), and functional assays showed that the increased transport activity of ZIP8 (C310G) was significantly suppressed by PPTD (Supplementary Fig. 6B), indicating the positive role of this pocket for metal transport and PPTD sensitivity.

**Fig. 6.**
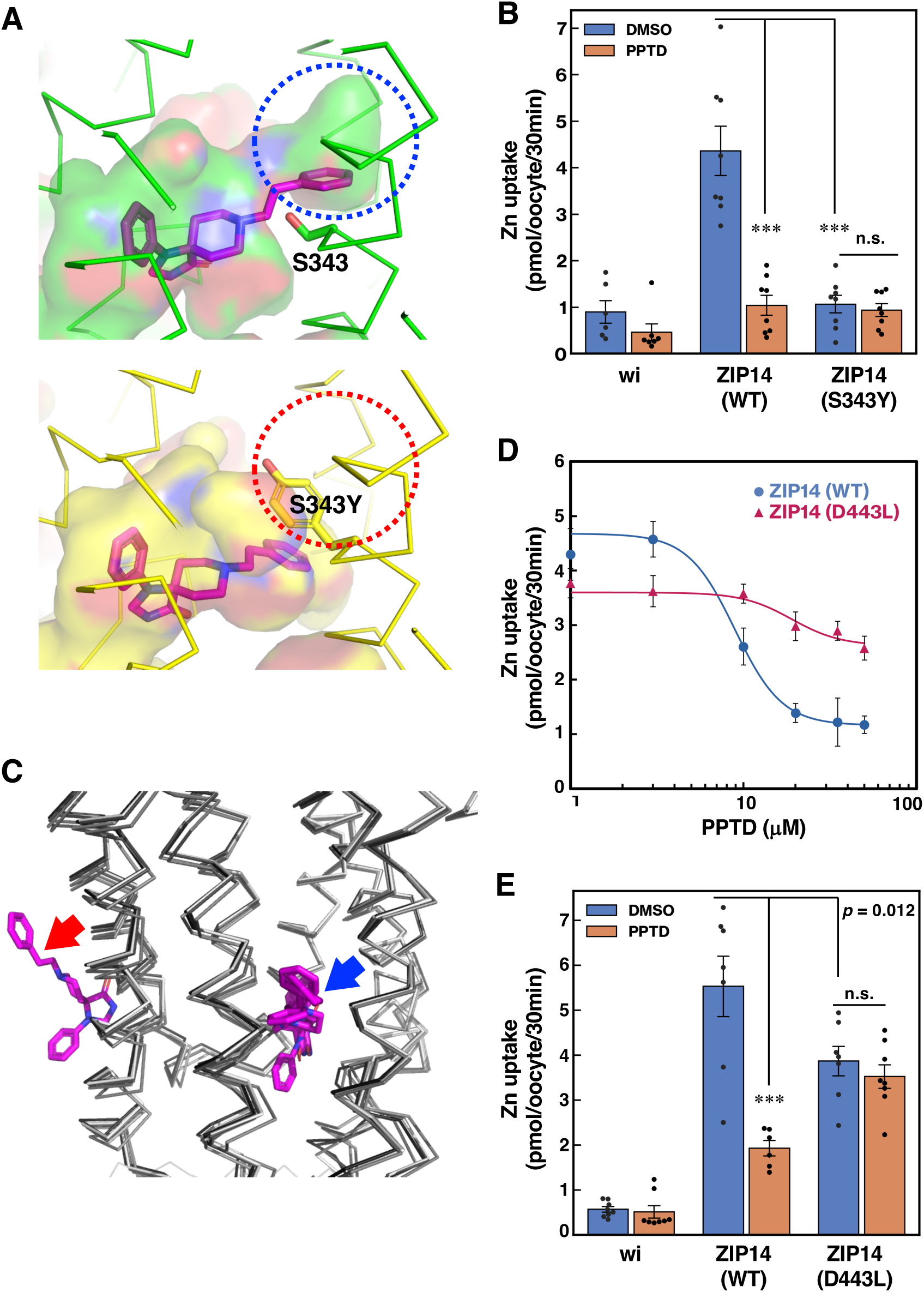
Identification of amino acids in ZIP14 required for the pocket formation and PPTD sensitivity. **A** Comparison of structural simulations of the human ZIP14 (WT)-PPTD complex (upper) and the ZIP14 (S343Y)-PPTD complex (lower). Substitution of Serine 343 with Tyrosine (S343Y) reduces the size of the PPTD-binding pocket within ZIP14, as indicated by the dotted red circle in the mutant structure. magenta: PPTD. **B** Serine 343 is essential for ZIP14-mediated zinc transport. *Xenopus* oocytes were injected either with human ZIP14 (WT), ZIP14 (S343Y) cRNA, or water (wi, control). The uptake assay was performed with 10 μM ZnSO_4_ including a trace amount of radioactive tracer in the presence of 50 μM PPTD. Radioactivity was measured as described under *Methods*. All data are shown as mean ± S.E.M. (n= 6 to 9 oocytes). **C** Structural overlay of five modeled complexes of PPTD bound to human ZIP14 with aspartic acid to leucine substitution at position 443 (D443L). In the four models, PPTD consistently occupies an internal binding region of ZIP14 (indicated by the blue arrow), while one model shows PPTD associating with the protein surface (the red arrow). This variation suggests that the D443L mutation may disrupt the stable binding of PPTD to ZIP14. **D** Aspartic acid at position 443 contributes to the sensitivity of ZIP14 to PPTD. After preincubation with indicated concentrations of PPTD, *Xenopus* oocytes injected with cRNA encoding wild-type human ZIP14 (WT) or the D443L mutant were incubated with 10 μM ZnSO_4_ including a trace amount of radioactive tracer in the presence of the same concentrations of PPTD for 30 min at 22 °C. The IC₅₀ values for ZIP14 WT and D443L were 8.9 μM and 19.0 μM, respectively. Radioactivity and IC₅₀ values were determined as described under *Methods*. **E** The ZIP14 D443L mutant retains transport activity but loses sensitivity to PPTD. *Xenopus* oocytes were injected with cRNA encoding either wild-type human ZIP14 (WT) or the D443L mutant. The uptake assay was performed with 10 μM ZnSO_4_ including a trace amount of radioactive tracer in the presence of 20 μM PPTD. Radioactivity was measured as described under *Methods*. Statistical analysis was performed using one-way ANOVA followed by Tukey’s multiple comparison test (\*\*\**p* < 0.001). All data are shown as mean ± S.E.M. (n= 6 to 9 oocytes).

In parallel, Asp443 contributed to PPTD stabilization via electrostatic interactions (Fig. 5C and Supplementary Table 5). Among the five AlphaFold3 docking models of the D443L mutant, in which Asp443 was substituted with leucine, one model showed a displacement of PPTD (Fig. 6C), suggesting reduced binding stability, which was confirmed by functional assays, with ZIP14 (D443L) mutant exhibiting a significantly elevated IC₅₀ (Fig. 6D) and reduced sensitivity to PPTD (Fig. 6E). Thus, Asp443 was suggested as another key determinant of PPTD’s efficacy. In addition, Phe234 and Leu346 also appear to form small hydrophobic subpockets that snugly accommodate the 2-phenylethyl chain of PPTD but lack sufficient space for bulkier substituent (Supplementary Movie 4 and 5).

These results delineate a detailed structural model for ZIP14-PPTD interaction, especially highlighting Ser343 and Asp443 of ZIP14 as essential for the binding pocket formation for PPTD and its sensitivity.

### PPTD ameliorates cachexia in a murine cancer model

PPTD inhibits mouse ZIP14-mediated metal transport in a manner similar to its inhibition on human ZIP14 (Supplementary Fig. 7). Given the role of ZIP14 in inflammatory conditions, including cancer-associated cachexia ^22^, we tested the therapeutic potential of PPTD, if any, in a mouse cachexia model (Fig. 7A). As expected, tumor-bearing mice exhibited progressive weight loss beginning approximately 20 days post-implantation (Fig. 7B left). PPTD administration tended to attenuate weight loss at both low and high doses (Fig. 7B middle and right, and Supplementary Fig. 8), and extended overall survival (Fig. 7C). Importantly, PPTD significantly delayed the progression of cachexia-related symptoms, including ≥10% body weight loss and death (Fig. 7D), prevented structural deterioration in the skeletal muscles and inflammation responses (Supplementary Fig. 9 and S10), and improved locomotor activity in the model (Fig. 7E and Supplementary Movie 7). These *in vivo* findings provide compelling evidence that PPTD can mitigate clinically important features of cachexia and highlight the therapeutic potential of targeting ZIP14 in inflammatory or wasting diseases (Fig. 7F).

**Fig. 7.**
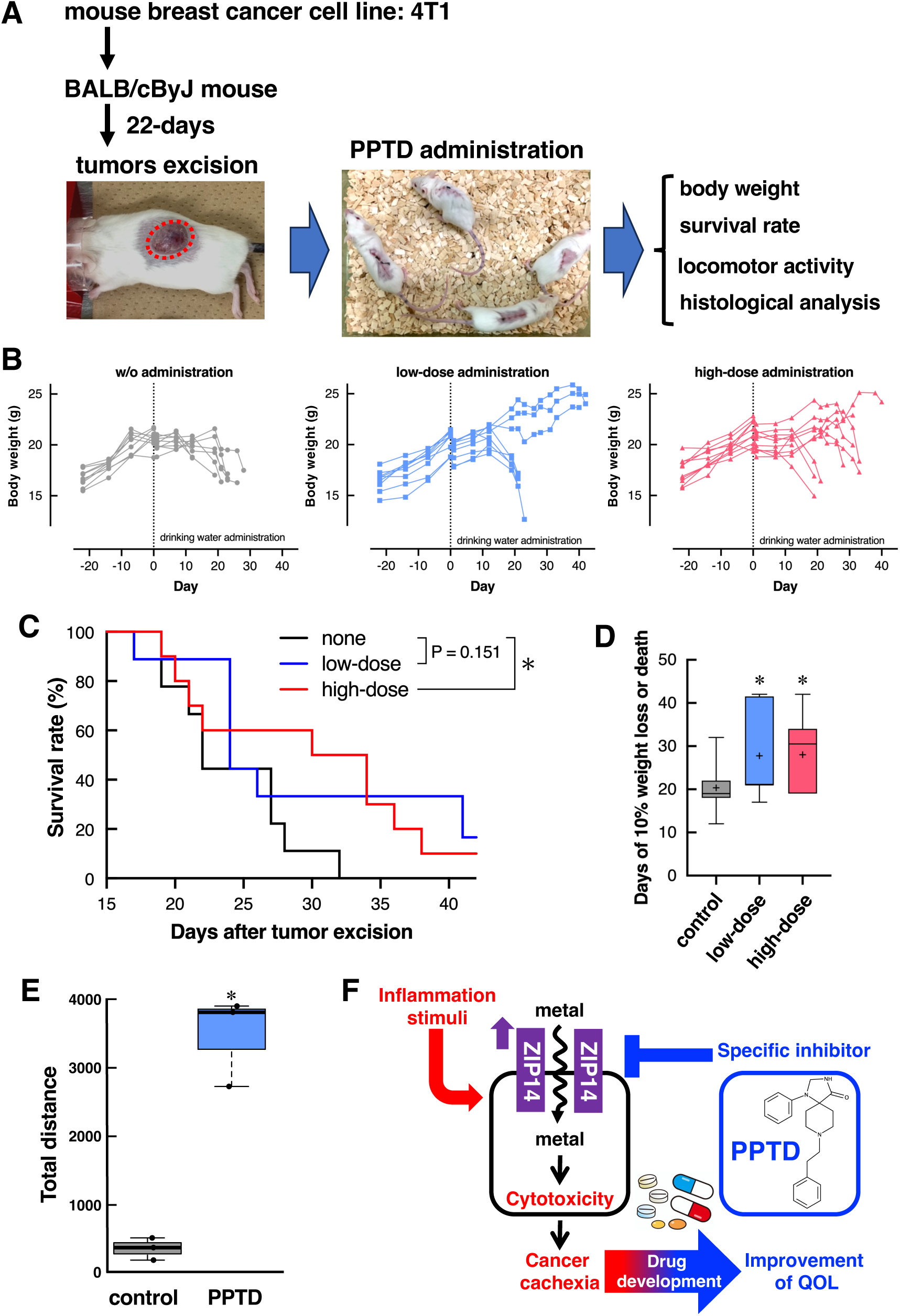
PPTD alleviates pathophysiological features in a mouse model of cancer cachexia. **A** Schematic overview of the experimental workflow for PPTD administration to the mouse model of cancer cachexia. **B** Effects of PPTD administration on body weight loss in the cancer cachexia model. Mice were administered PPTD via drinking water at either 0.1 g/L (low-dose) or 1.0 g/L (high-dose). **C** PPTD extends survival in the cancer cachexia model. Mice were administered 0.1 g/L (low dose, blue line) or 1.0 g/L (high dose, red line) PPTD through drinking water. Survival was analyzed using the Kaplan-Meier method. Black line: water-treated control group. Survival curves were compared using the Kaplan–Meier method and log-rank test (*p* < 0.05 is considered significant). **D** PPTD delays the onset of cachexia symptoms. Mice were administered 0.1 g/L (low-dose) or 1.0 g/L (high-dose) PPTD via drinking water. The day on which each group first exhibited ≥10% body weight loss or death is shown. Statistical analysis was performed using one-way ANOVA followed by Dunnett’s post hoc test (\**p* < 0.05). **E** PPTD improves locomotor activity in a cancer cachexia model. The locomotor of individual mice (n = 3 per group) was recorded on the 16th day of PPTD administration and analyzed using ImageJ (Supplementary Movie 7), as described under *Methods*. The results are presented as box plots with gray and blue boxes representing the H₂O control group and the PPTD-treated group (0.1 g/L), respectively. Each point represents an individual mouse. Statistical significance was assessed using Welch’s t-test to compare total distances between groups (\**p* < 0.05). **F** Working model: Inflammatory stimuli induce ZIP14 expression, promoting metal-induced cytotoxicity that contributes to cancer cachexia (left). The ZIP14 inhibitor PPTD alleviates key features of cancer cachexia (right) and may contribute to improving patients’ quality of life (QOL) (bottom).

## Discussion

ZIP14 has been implicated in various pathological conditions, largely due to its strong inducibility by inflammatory signals ^28–30^. Although ZIP14 shares high sequence homology and substrate specificity with its closest paralog, ZIP8 ^31^, their physiological roles diverge due to differences in tissue distribution and regulatory control. ZIP14 is highly expressed in the liver and pancreas ^18^, whereas ZIP8 is predominantly found in the lung, testis, and kidneys ^32^. Moreover, inflammatory stimuli robustly upregulate ZIP14 expression but only exert modest effects on ZIP8 ^28,33^. These functional and regulatory distinctions likely account for the varying clinical phenotypes associated with mutations in ZIP14 versus ZIP8 ^13,34^. The inducible nature of ZIP14 suggests that its excessive activity may contribute to disease pathogenesis in inflammation-associated conditions. Despite the growing recognition of ZIP14’s pathophysiological importance, no selective small-molecule modulators have previously been identified.

We identified PPTD as the first small-molecule inhibitor of ZIP14 by high-throughput screening. Originally synthesized as part of a structure–activity relationship study on μ-opioid receptor ligands ^35^, it exhibits weak inhibitory activity against glycine uptake via GlyT1 ^36^. However, to date, its broader biological functions or therapeutic applications have not been explored. Unlike previously reported ZIP7 inhibitors ^23^, PPTD possesses a structurally distinct scaffold. Although its selectivity for ZIP14 could not have been anticipated from existing SAR data, our docking simulations revealed that dimerized ZIP14 forms a unique internal pocket (Fig. 6A upper and Supplementary Fig. 5B, C) that can accommodate PPTD near the zinc-binding sites (Fig. 5B and Supplementary Movie 3). This pocket was absent in structural models of dimerized ZIP8 (Supplementary Fig. 5D), and the ZIP8 mutant with a newly formed pocket enhanced zinc transport and gained PPTD sensitivity (Supplementary Fig. 6), suggesting a structural basis for the observed selectivity and metal transport.

Dimerization of ZIP14 induced a conformational rearrangement wherein TM3 of one protomer could interact with TM4 of the opposing protomer (Supplementary Fig. 5A, right). This interaction remodels the TM4 region to create an internal pocket, forming a binding site for PPTD (Supplementary Fig. 5A, left panels). These findings are consistent with structural models suggesting that ZIP family dimerization, mediated by scaffold domains (TMs 2, 3, 7, and 8), stabilizes and modulates the transport domain (TMs 1, 4, 5, and 6), thereby facilitating substrate translocation ^37^. In particular, the interaction between TM3 of one protomer and TM4 of the opposing protomer plays a crucial role in forming a stable BMC ^27,38^. Notably, in the case of ZIP14, this structural configuration also gives rise to a unique PPTD-binding pocket at the dimer interface, a feature absent in ZIP8, thereby enabling isoform-specific inhibition. PPTD interacts with Ser343 (Fig. 5C and Supplementary Table 5), a residue involved in the formation of the PPTD-binding pocket (Fig. 6A), and in ZIP14-mediated metal transport (Fig. 6B), supporting the functional relevance of the pocket for PPTD sensitivity and ZIP14-mediated transport. Given that members of the ZIP family function as homo-or heterodimers ^3,5^, our findings may offer new insights into the rational design of compounds that can selectively regulate individual ZIP transporters.

PPTD was predicted to associate with His347 and His380 (Fig. 5C and Supplementary Table 5), which are part of conserved motifs in ZIP family members, namely HXXXE/D and HEXPHE, respectively. The HEXPHE motif contributes to BMC formation within the transport pathway and plays a key role in determining metal selectivity and facilitating transport. The HXXXE/D motif, located in TM4, serves as a metal-binding site and cooperates with the HEXPHE motif to coordinate BMC formation ^27^. Given that HXXXE/D and HEXPHE are key for interacting with metals for transport, their direct association with PPTD, giving rise to competitive inhibition of ZIP14-mediated transporting function is reasonable (Supplementary Fig. 11). These results provide a mechanistic rationale for the ZIP14 selectivity of PPTD and highlight a novel mode of transporter inhibition via ligand recognition at a dimerization-induced interface.

What makes ZIP14 a notable potential drug target? Unlike other closely related ZIP transporters, ZIP14 is a unique inducible transporter—one whose expression is dramatically upregulated by inflammatory signals, such as LPS, IL-1β, IL-6, and nitric oxide ^39^. ZIP14 itself enhances inflammatory signaling through a positive feedback loop ^30^. Therefore, ZIP14 is a mediator and a potential amplifier of inflammatory damage, positioning ZIP14 at the crossroads of metal homeostasis and inflammation-related pathology and making it an attractive target for therapeutic intervention, including cancer cachexia ^22^, a syndrome characterized by systemic inflammation, muscle wasting, and metabolic dysregulation ^40^. Despite advances in identifying inflammatory cytokines associated with cancer cachexia, effective clinical treatments remain elusive. According to the international consensus led by the European Palliative Care Research Collaborative (EPCRC), cancer cachexia is defined as a multifactorial syndrome that cannot be fully reversed by conventional nutritional support. The consensus further classifies the condition into stages—pre-cachexia, cachexia, and refractory cachexia—and emphasizes the importance of comprehensive, multidisciplinary intervention beginning in the early stages (i.e., pre-cachexia or cachexia) ^41^. PPTD may serve as a novel therapeutic agent for ZIP14-induced cancer cachexia at early stages, as it delayed disease progression (Fig. 7D), prevented skeletal muscle deterioration (Supplementary Fig. 9), and consequently improved locomotor activity (Fig. 7E and Supplementary Movie 7). Given that PPTD attenuated key clinical features of cachexia in the model, our findings underscore its potential not only as part of a combinatorial therapeutic strategy for cancer cachexia but also as a means to enhance quality of life (QOL) and maintain patient dignity during the course of illness (Fig. 7F).

ZIP14 has been implicated in iron overload disorders ^21^, where it facilitates hepatic uptake of NTBI, a redox-active form of iron that promotes oxidative stress through ROS and LPO generation, ultimately leading to iron toxicity ^42^. Current treatments rely on phlebotomy or iron chelators ^43^; however, novel strategies targeting NTBI uptake pathways are needed. Our findings demonstrate that PPTD suppresses ZIP14-mediated ROS and LPO production (Fig. 4) and inflammatory response (Supplementary Fig. 10), underscoring its potential as a therapeutic scaffold for iron overload-related conditions.

Our findings establish PPTD as the first small-molecule compound that specifically inhibits ZIP14. PPTD is a promising lead compound for probing the biological functions of ZIP14 and for developing therapeutics targeting this druggable metal transporter.

## Methods

### Transfection, cell lines, immunoblotting, and morphological analysis

HEK293 cells and the C2C12 mouse myoblast cells were cultured in Dulbecco’s Modified Eagle Medium (DMEM) supplemented with 10% fetal bovine serum (FBS) and antibiotics. Lipofectamine™ 2000 (Thermo Fisher Scientific, #11668019) was used to transfect the human ZIP14 plasmid into HEK293 or C2C12 cells following the manufacturer’s instructions. To generate inducible cell lines, the coding sequences (CDS) of human ZIP14 (NM_015359.1) and human ZIP8 (NM_001135146) were inserted into the pcDNA4/TO mammalian expression vector (Invitrogen). These constructs were used to establish HEK293 cell lines that express hZIP14 and human ZIP8 under the control of a tetracycline-regulated (T-REx) system, following the manufacturer’s instructions. For inducible expression, cells were treated with 1 μg/mL Tet in DMEM supplemented with 10% FBS for the indicated durations. Cells were lysed in buffer supplemented with protease inhibitors (Nacalai Tesque, #08714-04), and clear lysates were obtained by centrifugation at 10,000 × *g* for 10 min at 4 °C. Equal amounts of protein (20 μg) from each sample were resolved by SDS–PAGE and subjected to western blotting (Supplementary Fig. 1). The following primary antibodies were used: anti-human ZIP14 (generated previously ^12^), anti-human ZIP8 (Alomone Labs, #AZT-008), and anti-β-actin (Cell Signaling Technology, #3700S). Cell morphology was assessed using phase-contrast microscopy on the BIOREVO BZ-9000 microscope system (Keyence).

### Screening inhibitors of human ZIP14-mediated zinc uptake

We established HEK 293 cell lines with inducible expression of human ZIP14 or ZIP8 through Tet induction to identify chemical compounds that inhibit zinc uptake mediated by human ZIP14 (TREx-hZIP14 and hZIP8) (Supplementary Fig. 1A, B). These cells were utilized for both primary, secondary, and tertiary screens (Fig. 1A). For screening, cells were cultured in DMEM supplemented with 10% low-zinc FBS, prepared by incubating FBS with Chelex 100 resin (BioRad) for 15 h to deplete zinc, and treated with 1 μg/mL of Tet for 24 h. Following Tet induction, the cells were resuspended in PBS containing 2% low-zinc FBS and pre-incubated for 30 min with 1 μM of the fluorescent intracellular zinc indicator FluoZin-3 AM (Thermo Fisher Scientific), 1 μg/mL of Hoechst 33342 (Thermo Fisher Scientific), and 10 μM of each compound from a proprietary chemical library containing approximately 30,000 small molecules, or DMSO as a control. Cells were incubated with 5 μM ZnSO₄ for 1 h, and intracellular zinc levels were quantified using the IN Cell Analyzer 6000 (GE Healthcare).

### Transport assay in *Xenopus* oocytes

The inhibitory activities of the candidate compounds and their analogs were evaluated using a metal transport assay in *Xenopus laevis* oocytes, as previously described with minor modifications ^44^. HA-tagged human ZIP14 isoform 1 and ZIP8 isoform 1 cDNAs were amplified by PCR and cloned into the pcDNA3.1(-) mammalian expression vector into the EcoRV site using the In-Fusion HD Cloning Kit (Takara Bio) following the manufacturer’s instructions. The primers contained an HA-tag sequence and 15 bp overlapping regions with the vector ends. Mutations of ZIP14 were introduced by PCR using Q5 High-Fidelity DNA Polymerase (New England Biolabs) and In-Fusion HD Cloning Kit. All primer sequences are available upon request. The resulting plasmids were linearized with *HindIII*, and capped cRNAs were transcribed in vitro using the mMESSAGE mMACHINE T7 Transcription Kit (Thermo Fisher Scientific). Transcripts were polyadenylated and purified using the Poly(A) Tailing Kit and the MEGAclear Transcription Clean-Up Kit (Thermo Fisher Scientific) following the manufacturer’s protocols.

To prepare oocytes, stage IV–VI *Xenopus* oocytes were harvested and enzymatically defolliculated with 1.0–1.5 mg/mL collagenase A (Roche) in OR2 buffer (82 mM NaCl, 2.5 mM KCl, 1 mM MgCl₂, 5 mM HEPES, pH 7.6) for 1.0–1.5 h at 25 °C. After washing, oocytes were transferred to ND96 buffer (96 mM NaCl, 2 mM KCl, 1.8 mM CaCl₂, 1 mM MgCl₂, 5 mM HEPES, pH 7.5), and injected with 23 nL (23 ng) of purified cRNA or RNA-free water using a Nanoject II microinjector (Drummond Scientific). Oocytes were incubated at 18 °C for 2 days in ND96 buffer to facilitate transporter expression. Expression of HA-tagged human ZIP14 and human ZIP8 was verified using immunofluorescence microscopy (Supplementary Fig. 3). Oocytes were fixed with 4% paraformaldehyde, embedded in paraffin, and stained with an anti-HA antibody (Covance, MMS-101R; 1:50) followed by the corresponding Alexa Fluor 488-conjugated anti-mouse IgG secondary antibody (Thermo Fisher Scientific; 1:100). Fluorescent and differential interference contrast (DIC) images were acquired using a confocal laser scanning microscope (Fluoview FV1000; Olympus).

For the metal uptake assay, transporter-expressing oocytes were washed twice and preincubated in 96 well plate for 30 min in ND96 buffer containing either PPTD or DMSO (control). For iron uptake, oocytes were incubated in ND96 buffer supplemented with 10 μM FeCl₂, 5 μCi/mL [⁵⁵Fe]Cl₃ (Perkin Elmer), and 1 mM ascorbic acid for 30 min at 22 °C. Following incubation, oocytes were transferred to 24 well plate and washed five times with ice-cold ND96 buffer containing 1 mM ascorbic acid and lysed in 0.0625 N NaOH. Radioactivity was quantified using liquid scintillation counting. For zinc uptake, oocytes were incubated in 96 well plate with ND96 buffer containing 10 μM ZnSO_4_ and 2 μCi/mL [⁶⁵Zn]Cl₂ (RIKEN) unless otherwise stated. After five washes with ice-cold ND96 containing 5 mM EDTA in 24 well plate, uptake was visualized using a phosphorimager (Typhoon FLA 9500; GE Healthcare) and quantified using the ImageJ software. Data are presented as mean ± S.E.M with individual oocyte values shown as dots (n= 6 to 9 oocytes) unless otherwise stated. Reproducibility of the results was confirmed by 2-3 independent experiments using different batches of oocytes. Statistical significance was determined using one-way ANOVA followed by Tukey’s post hoc test.

### Cytotoxicity assays

For cytotoxicity assessment, TREx-hZIP14 and TREx-hZIP8 cells (2 × 10⁵ cells per well in six-well plates) were cultured in Tet-containing DMEM at 37 °C in an atmosphere with 5% CO₂ for 24 h to induce transgene expression. Cells were incubated for an additional 24 h with the indicated concentrations of PPTD or its analogs in the presence of divalent metal ions: Zn²⁺ (100 μM), Fe²⁺ (20 μM), Mn²⁺ (100 μM), or Cd²⁺ (1 μM). Cytotoxicity mitigation by PPTD was evaluated using phase-contrast microscopy (for zinc and iron treatments) and cell viability using the alamarBlue assay (Invitrogen) with a microplate reader (for manganese and cadmium treatments). To analyze Mn and Cd uptake, TREx-hZIP14 and TREx-hZIP8 cells (2 × 10⁵ cells per well in six-well plates) were similarly preincubated with a Tet-containing medium for 24 h. Cells were treated with the indicated concentrations of PPTD in the presence of radiolabeled metal ions, ⁵⁴Mn (0.1 μM) or ¹⁰⁹Cd (0.1 μM), for 1 h. Subsequently, cells were washed with an FBS-containing medium, followed by PBS-EDTA to remove extracellular metal ions. The amount of intracellularly accumulated metal was quantified using a γ-counter (AccuFLEX γ7000; Hitachi).

### Assessment of ROS and LPO generation

TREx-hZIP14 or TREx-hZIP8 cells were cultured in the presence of Tet (1 µg/mL) to induce transporter expression. The cells were treated with or without 10 µM PPTD for the indicated periods. Following treatment, intracellular ROS and LPO levels were measured using CellROX™ Green Reagent (Thermo Fisher Scientific) and Liperfluo™ (Dojindo), respectively. The cells were incubated with 5 µM CellROX Green or Liperfluo™ in culture medium for 30 min at 37 °C in a humidified atmosphere with 5% CO₂. Subsequently, the cells were harvested, resuspended in PBS supplemented with 2% FBS, and the fluorescence intensity, reflecting intracellular ROS or LPO generation, was analyzed by flow cytometry (Guava system, Millipore) and using the BIOREVO BZ-9000 microscope system (Keyence).

### Organization of PPTD and its derivatives

The synthesis methods and NMR data for all PPTD derivatives (Supplementary Tables 2, 3, and 4) are provided under *Supplementary Information*.

### Computational modeling of PPTD and ZIP14 complex

Locally installed AlphaFold3 (version 3.0.0) was used to model the structure of PPTD and human ZIP14 (or ZIP8) complex ^45^. The amino acid sequence of human ZIP14 and the chemical structure of PPTD, represented in SMILES format, were included in an input JSON file for the prediction. Five predicted models were analyzed using a locally installed PLIP (version 2.4.0) ^46^ to identify interacting residues. All molecular images and movies were generated using PyMOL (version 2.5.0). The pockets were visualized using the surface representation with the “Cavities and Pockets (Culled)” setting in PyMOL.

### Animal experiments

All animal experiments were conducted following the ARRIVE guidelines and the Tokushima-Bunri University, the Kyushu University Animal Experimentation Regulations, and were approved by the Institutional Animal Care and Use Committee of Kyushu University (Approval Number: A24-231-0; Approval Date: March 26, 2024) and of Tokushima Bunri University (Approval Number 19-2; Approval Date: April 1, 2024), respectively.

#### Establishing the mouse model of cancer cachexia

A murine model of cancer cachexia was established as previously described ^22^. Briefly, four-week-old female BALB/cByJ mice (Jackson Laboratory Japan, Kanagawa) were acclimated for one week before initiating experiments at five weeks of age. Mice were housed under standard conditions (24 ± 1°C, 60% ± 10% humidity, 12-h light/dark cycle) with free access to food (CE-2, CLEA Japan) and water. 4T1 breast cancer cells (RRID: CVCL_0125; RIKEN BioResource Center) were cultured in RPMI-1640 medium supplemented with 5% FBS and 0.25% penicillin-streptomycin at 37 °C in 5% CO₂. A total of 1 × 10⁵ cells in 200 μL of 1:1 Matrigel/PBS were subcutaneously injected into the dorsal region of isoflurane-anesthetized mice.

#### Oral administration of PPTD-HCl via drinking water

Starting from Day 0 post-tumor resection, mice were administered PPTD-HCl via drinking water. PPTD-HCl was dissolved in tap water at concentrations of 0.1 and 1.0 g/L and supplied in water bottles, which were replaced twice weekly. From Day 0 to Day 7, the water intake of each group was measured. Body weight was recorded throughout the experimental period, and survival was monitored daily following tumor resection. From Day 10 onward, several mice exhibited deterioration in health, making access to the standard feed located at the top of the cage difficult. To accommodate these animals, feed was additionally placed at each of the four corners of the cage floor.

#### Histological and locomotor activity analysis

Histological analysis of skeletal muscle samples was performed as previously described ^47^. The locomotor of individual mice (n = 3 per group) was recorded on the 16th day of PPTD (0.1 g/L) administration (Supplementary Movie 7), and analyzed using ImageJ. Head-centered tracking was manually performed frame by frame across 46 time points to determine locomotor trajectories. Total travel distances were calculated based on changes in X and Y coordinates between successive frames and expressed in pixels, which are presented as box plots (Fig. 7E).

## Supporting information

Supplementaryl informaiton

## Acknowledgements

This study was supported by grants from the Japan Society for the Promotion of Science (JSPS) KAKENHI (17H04011, 20H03409 and 25K09964 to T.F.; 21J40212, 22K11871 and 22KJ3069 to E.Y.), the Astellas Foundation for Research on Metabolic Disorders, Astellas Pharma Inc., the Mitsubishi Foundation, the Naito Foundation, the Princess Takamatsu Cancer Research Fund, the Science Research Promotion Fund, Takeda Science Foundation, the Uehara Memorial Foundation, the Vehicle Racing Commemorative Foundation, and the Yasuda Medical Foundation (to T.F.), and the US National Institutes of Health grant R01 DK-080706 (to M.D.K).

This research was also supported by Research Support Project for Life Science and Drug Discovery (Basis for Supporting Innovative Drug Discovery and Life Science Research: BINDS) from AMED under Grant Number JP25ama121054 and JP25ama121052. We thank Dr. Keiichiro Uchida and Dr. Rong Cheng (Kyoto University) for their contributions to chemical synthesis.

## Ethics declarations Competing Interests

This research was supported in part by Astellas Pharma Inc.

