## Supplementaryl informaiton for "Discovery of a Selective Inhibitor of ZIP14 with Therapeutic Potential for Cancer-associated Cachexia"

**This PDF file includes:**

Supplementary Methods: page S2-S23

Supplementary Figs 1 to 11: page S24-S34

Supplementary Tables 1 to 6: page S35-S40

Supplementary Reference: page S41

Legends for Supplementary Movies: page S42

**Other supplementary materials for this manuscript include the following:**

Supplementary Movies 1 to 7

#### Supplementary Methods

#### Chemical synthesis of PPTD

All reactions sensitive to moisture and/or air were conducted in an atmosphere of argon in dry solvents under anhydrous conditions using oven-dried glassware unless otherwise stated. Anhydrous dichloromethane ($CH_2Cl_2$) and *N*,*N*-dimethylformamide (DMF) were purchased from Kanto Chemical Co. Inc. and used directly without further drying. 1-Phenyl-1,3,8-triazaspiro[4.5]decan-4-one (PTD) was purchased from FUJIFILM Wako Pure Chemical Co. and used directly without purification. All other chemicals of the highest commercial grade were purchased and directly used. Analytical thin-layer chromatography (TLC) was performed using E. Merck silica gel 60 $F_{254}$ plates (0.25 mm thickness). Column chromatography was performed using Kanto Chemical silica gel 60N (40–100 mesh, spherical, neutral). IR spectra were recorded on a JASCO FT/IR-4200 spectrometer. $^{1}$H and $^{13}$C NMR spectra were recorded on a Bruker ADVANCE III HD spectrometer (500 MHz) at room temperature unless otherwise stated. Chemical shift values are reported in ppm (δ) downfield from tetramethylsilane against the internal solvent [$^{1}$H NMR, C*H*$Cl_3$ (7.26), C*H*$D_2$OD (3.31); $^{13}$C NMR, *C*$DCl_3$ (77.0), *C*$D_3$OD (49.0)] unless otherwise indicated. Coupling constants (*J*) are reported in hertz (Hz). The following abbreviations were used to designate the multiplicities: s = singlet; d = doublet; t = triplet; m = multiplet; br = broad. CI mass spectra were measured on a JEOL MStation JMS-700.

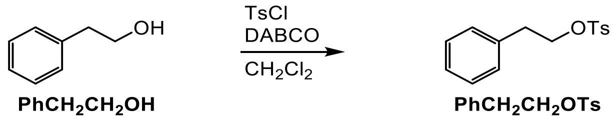


**2-phenylethyl 4-methylbenzenesulfonate.** To a solution of $PhCH_2CH_2OH$ (1.956 g, 16.012 mmol), DABCO (3.599 g, 32.082 mmol) in $CH_2Cl_2$ (80 mL), at room temperature, TsCl was added (4.575 g, 23.998 mmol). After stirring at room temperature for 3 d, the resulting solution containing white precipitates was passed through Celite, washed with EtOAc, and the filtrate was concentrated under reduced pressure. The resultant oil was dissolved in EtOAc, and the organic layer was washed with sat. aq $NaHCO_3$, brine, dried over $Na_2SO_4$, filtered, and concentrated under reduced pressure. Purification of the residue by column chromatography (silica gel, hexanes/EtOAc = 20/1 to 2/1) yielded $PhCH_2CH_2OTs$ (4.140 g, 14.981 mmol, 94%) as a colorless oil: IR (film) 3029, 2959, 1598, 1496, 1455, 1354, 1188, 1173, 1096, 960, 915, 890, 814, 774, 752, 698, 661, 552 $cm^{-1}$; $^{1}$H NMR (500 MHz, $CDCl_3$) δ 7.68 (m, 2H), 7.28–7.18 (m, 5H), 7.10 (m, 2H), 4.20 (t, *J* = 7.0 Hz, 2H), 2.94 (t, *J* = 7.0 Hz, 2H), 2.42 (s, 3H); $^{13}$C NMR (125 MHz, $CDCl_3$) δ 144.6, 136.1, 132.9, 129.7 (2C), 128.8 (2C), 128.5 (2C), 127.7 (2C), 126.8, 70.5, 35.3, 21.5; HRMS (CI) calcd for $C_{15}H_{17}O_3SNa$ [$(M+H)^+$] 277.0893, found 277.0897.

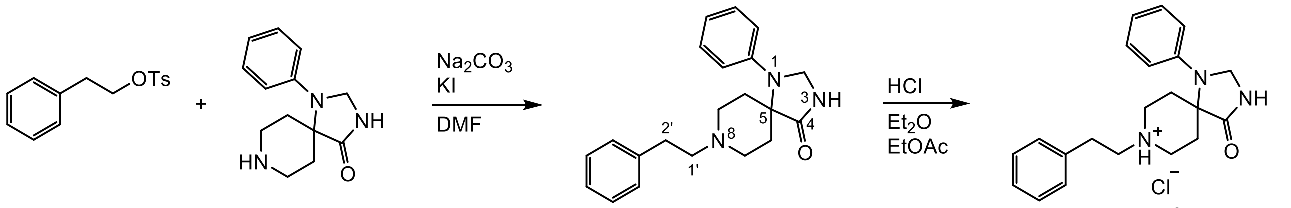


**PhCH$_2$CH$_2$OTs** **PTD** **PPTD** **PPTD·HCl**

**4-oxo-8-phenethyl-1-phenyl-1,3,8-triazaspiro[4.5]decan-8-ium chloride (**PPTD·HCl**).** A solution of $PhCH_2CH_2OTs$ (3.827 g, 13.848 mmol), PTD (3.051 g, 13.189 mmol), $Na_2CO_3$ (2.8 g, 26.417 mmol), KI (232.6 mg, 1.401 mmol) in DMF (27.5 mL) was stirred at 80 °C for 14 h, and subsequently $H_2O$ (~30 mL) was added at room temperature, resulting in the formation of a white solid. The solid was collected by filtration, washed with $H_2O$ and dried under reduced pressure to obtain PPTD (3.730 g, 11.119 mmol, 84%) as a white solid: IR (film) 3209, 2926, 2826, 2560, 1704, 1598, 1499, 1372, 748, 697 cm$^{-1}$; $^1$H NMR (500 MHz, $CD_3OD$) δ 7.34–7.28 (m, 6H, $H^{Ph}$), 7.24 (m, 1H, $H^{2'-Ph}$), 7.05 (m, 2H, $H^{N\text{-}Ph}$), 6.92 (m, 1H, $H^{N\text{-}Ph}$), 4.73 (s, 2H, $H^2$), 3.60 (m, 2H, $H^{7ax}$), 3.42 (m, 2H, $H^{7eq}$), 3.17 (m, 2H, $H^{1'}$), 3.03 (m, 2H, $H^{2'}$), 2.75 (m, 2H, $H^{6ax}$), 1.97 (br d, $J$ = 15.0 Hz, 2H, $H^{6eq}$) (1H was missing due to H–D exchange); $^{13}$C NMR (125 MHz, $CD_3OD$) δ 177.9 ($C^4$), 144.2 ($C^{Ph}$), 138.7 ($C^{Ph}$), 130.4 (2C, $C^{Ph}$), 129.9 (2C, $C^{Ph}$), 129.8 (2C, $C^{Ph}$), 128.0 ($C^{Ph}$), 121.5 ($C^{Ph}$), 118.4 (2C, $C^{Ph}$), 60.8 ($C^2$), 59.8 ($C^{1'}$), 59.4 ($C^5$), 49.3 (2C, $C^7$), 32.3 ($C^{2'}$), 28.9 (2C, $C^6$); HRMS (CI) calcd for $C_{21}H_{26}N_3O$ [(M+H)$^+$] 336.2070, found 336.2079.

PPTD was dissolved in hot EtOAc, and to the solution, 1 M HCl in $Et_2O$ (15 mL) was added, resulting in the formation of a white precipitate immediately. The solid was filtered, washed with $H_2O$, dried in vacuo, and recrystallized from EtOH, yielding PPTD·HCl (4.0193 g, 10.807 mmol) as a white solid: IR (film) 3397, 3201, 2936, 2565, 1705, 1599, 1498, 1373, 751, 698 cm$^{-1}$; $^1$H NMR (500 MHz, $CD_3OD$) δ 7.36–7.31 (m, 6H), 7.27 (m, 1H), 7.07 (m, 2H), 6.95 (m, 1H), 4.75 (s, 2H), 3.87 (m, 2H), 3.63 (m, 2H), 3.39 (m, 2H), 3.12 (m, 2H), 2.82 (m, 2H), 2.07 (d, $J$ = 15.3 Hz, 2H) (2H was missing due to H–D exchange); $^{13}$C NMR (125 MHz, $CD_3OD$) δ 177.4, 143.9, 137.6, 130.5 (2C), 130.0 (2C), 129.8 (2C), 128.3 (2C), 121.7, 118.5, 60.8, 59.2, 58.8, 50.6 (2C), 31.5 (2C), 28.5.

**X-ray crystallography**

Single crystals of PPTD, obtained by slow evaporation of hexanes/ethyl acetate, were selected and fitted onto a glass fiber and subjected to measurement at −173 °C with a Bruker Apex II ultra diffractometer using MoKα radiation. Data correction and reduction were performed with the crystallographic package Apex 3. The structure was solved using the intrinsic phasing method and refined by full matrix least-squares method based on $F^2$ using SHELXL-2014/7 (Sheldrick, 2014). All non-hydrogen atoms were refined anisotropically, and hydrogen atoms were geometrically positioned. A total of 901 parameters were finally considered. Final disagreement indices were R1 = 0.0357, wR2 = 0.1003 (I > 2 sigma (I)). The ORTEP plot was obtained using PLATON (A. L. Spek, 2009).

Crystal data: colorless needle, $C_{21}H_{25}N_3O$, MW = 335.44, monoclinic, space group *C1c1*, $Z$ = 16, $a$ = 27.183(3) Å, $b$ = 12.2338(11) Å, $c$ = 20.9462(19) Å, $\beta$ = 94.254(2) °, $V$ = 6946.5(11) Å$^3$, GOF = 1.035.

Crystallographic data for the structures of PPTD have been deposited with the Cambridge Crystallographic Data Centre under supplementary publication number CCDC 2456161. Copies of the data can be obtained, free of charge, on application to CCDC, 12 Union Road, Cambridge CB2 1EZ UK (Fax: +44(0)-1223-336033 or).

Single crystals of PPTD·HCl obtained by slow evaporation of MeOH, were selected and fitted onto a glass fiber and subjected to measurement at −173 °C on a Bruker Apex II ultra diffractometer using MoKα radiation. Data correction and reduction were performed using the crystallographic package Apex 3. The structure was solved by the intrinsic phasing method and refined by full matrix least-squares method based on $F^2$ using SHELXL-2014/7 (Sheldrick, 2014). All non-hydrogen atoms were anisotropically refined, and all hydrogen atoms except for one acidic hydrogen (difmap) on the tertiary amine were positioned geometrically. A total of 477 parameters were finally considered. Final disagreement indices were R1 = 0.0358, wR2 = 0.1062 (I > 2 sigma (I)). The ORTEP plot was obtained using PLATON (A. L. Spek, 2009).
Crystal data: colorless cube, $C_{21}H_{26}ClN_3O$, MW = 371.90, Orthorhombic, space group $Pca2_1$, $Z$ = 8, $a$ = 10.005(7) Å, $b$ = 9.973(7) Å, $c$ = 39.52(3) Å, $V$ = 3943.(5) Å$^3$, GOF = 1.053.
Crystallographic data for the structures of PPTD·HCl have been deposited with the Cambridge Crystallographic Data Centre under the supplementary publication number CCDC 2456157. Copies of the data can be obtained, free of charge, on application to CCDC, 12 Union Road, Cambridge CB2 1EZ UK (Fax: +44(0)-1223-336033 or).

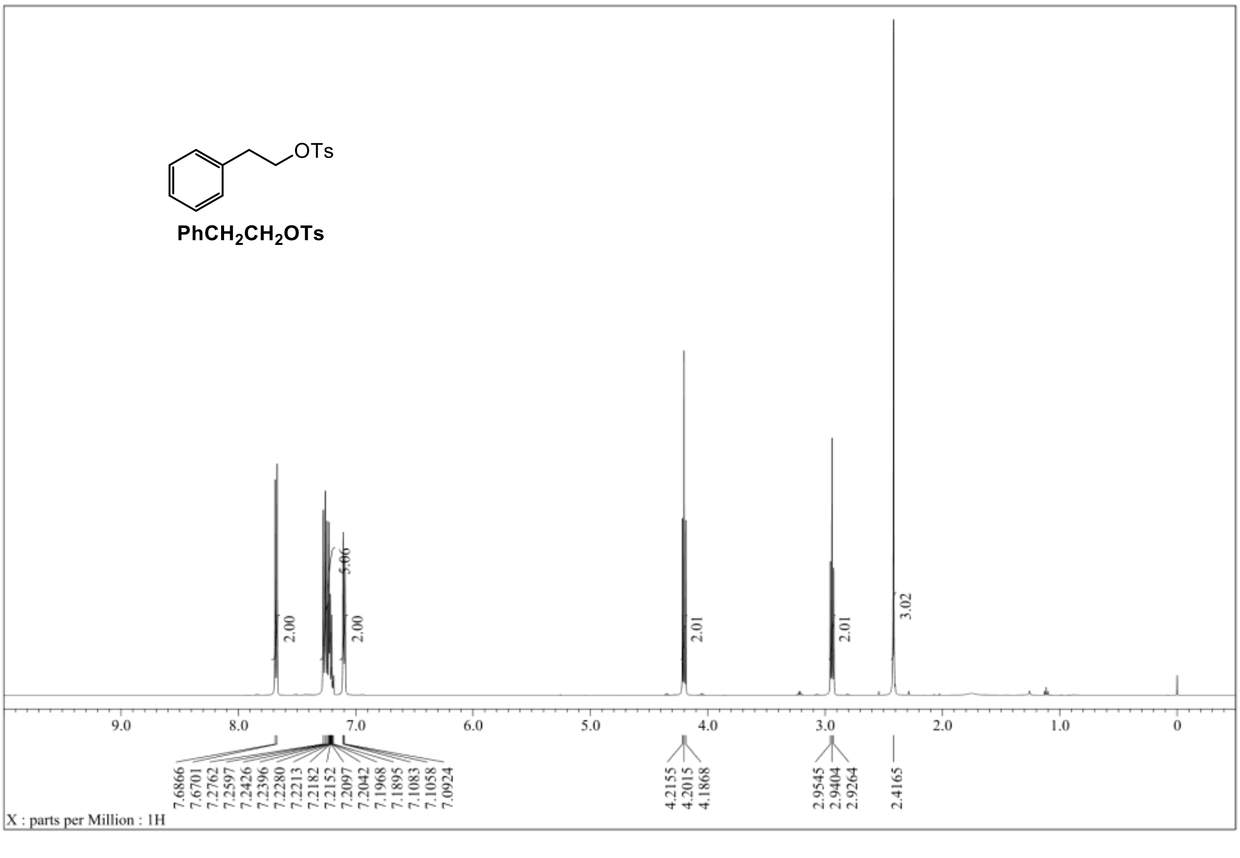


$^1$H NMR spectrum of $PhCH_2CH_2OTs$ (500 MHz, $CDCl_3$).

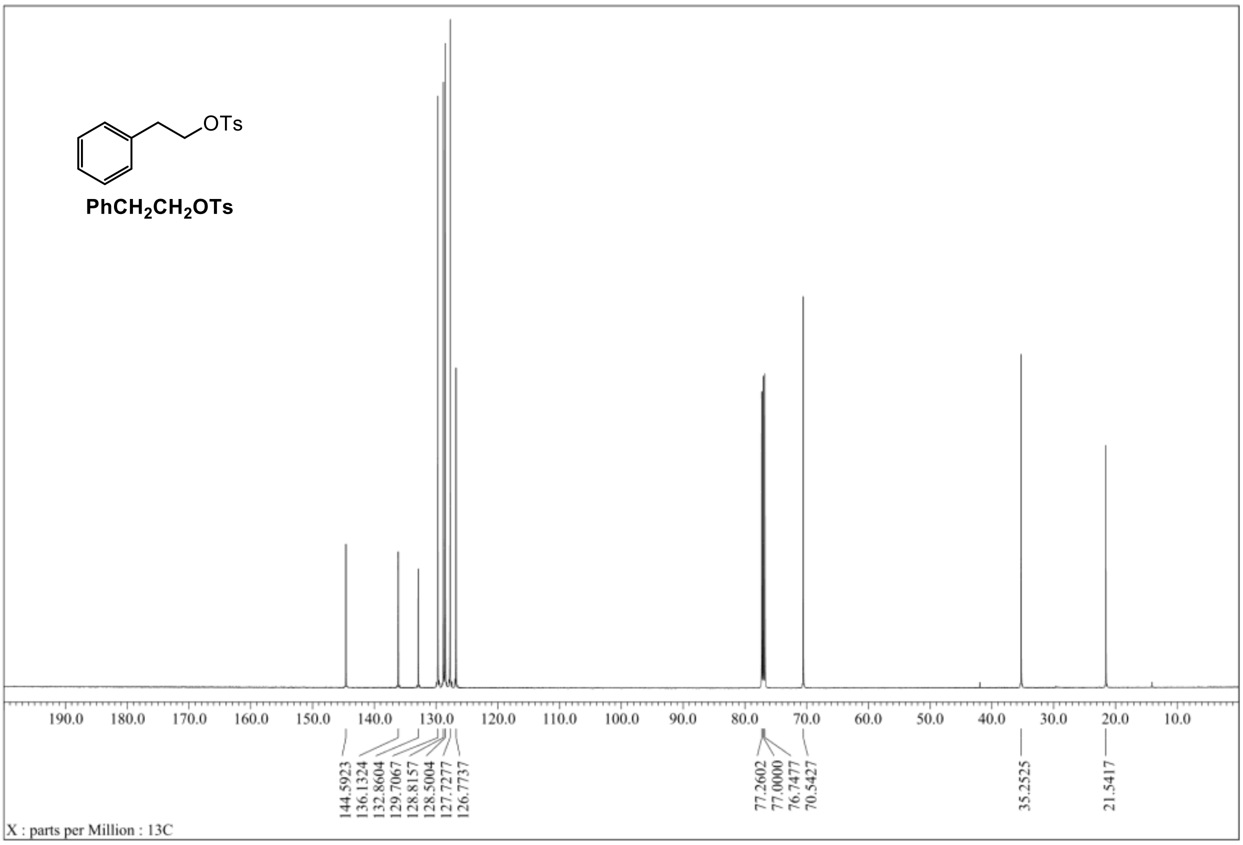


$^{13}C$ NMR spectrum of $PhCH_2CH_2OTs$ (125 MHz, $CDCl_3$).

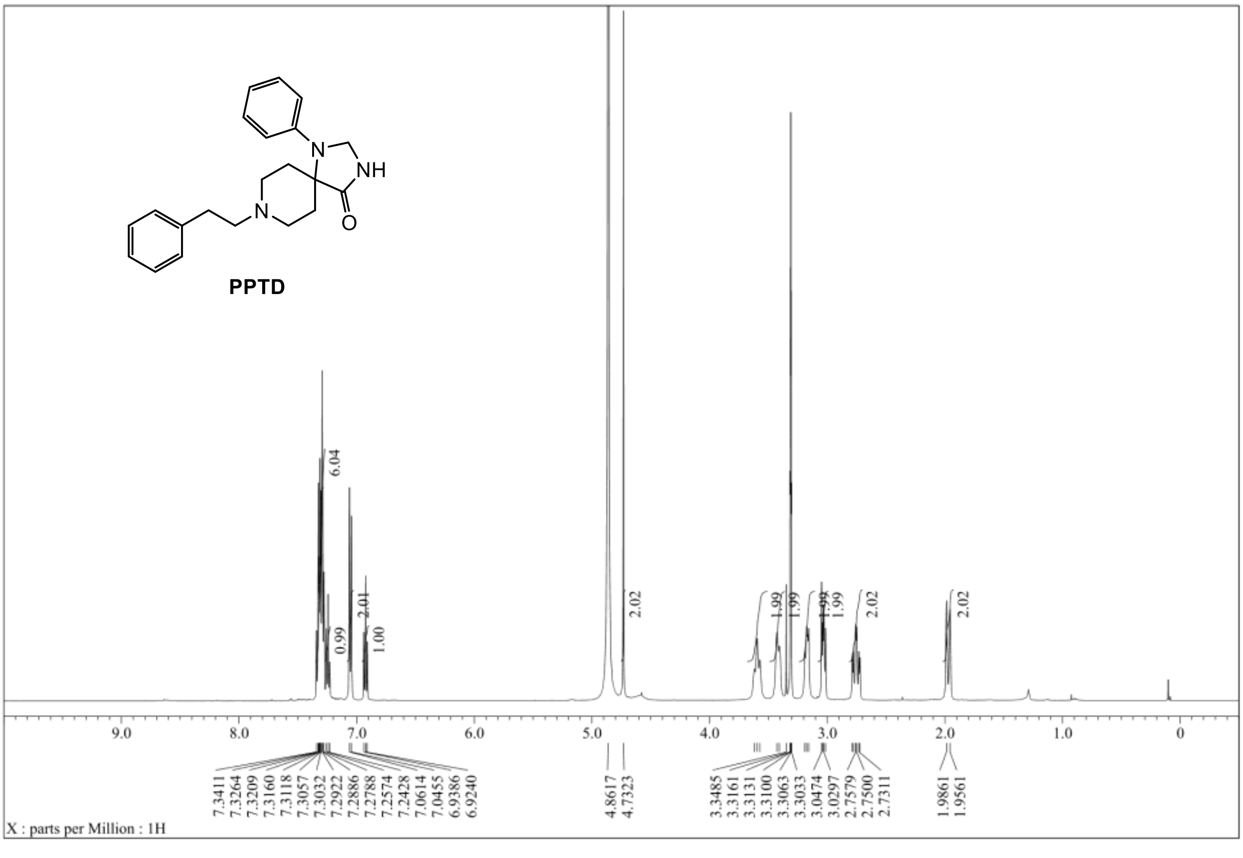


$^{1}H$ NMR spectrum of PPTD (500 MHz, $CD_3OD$).

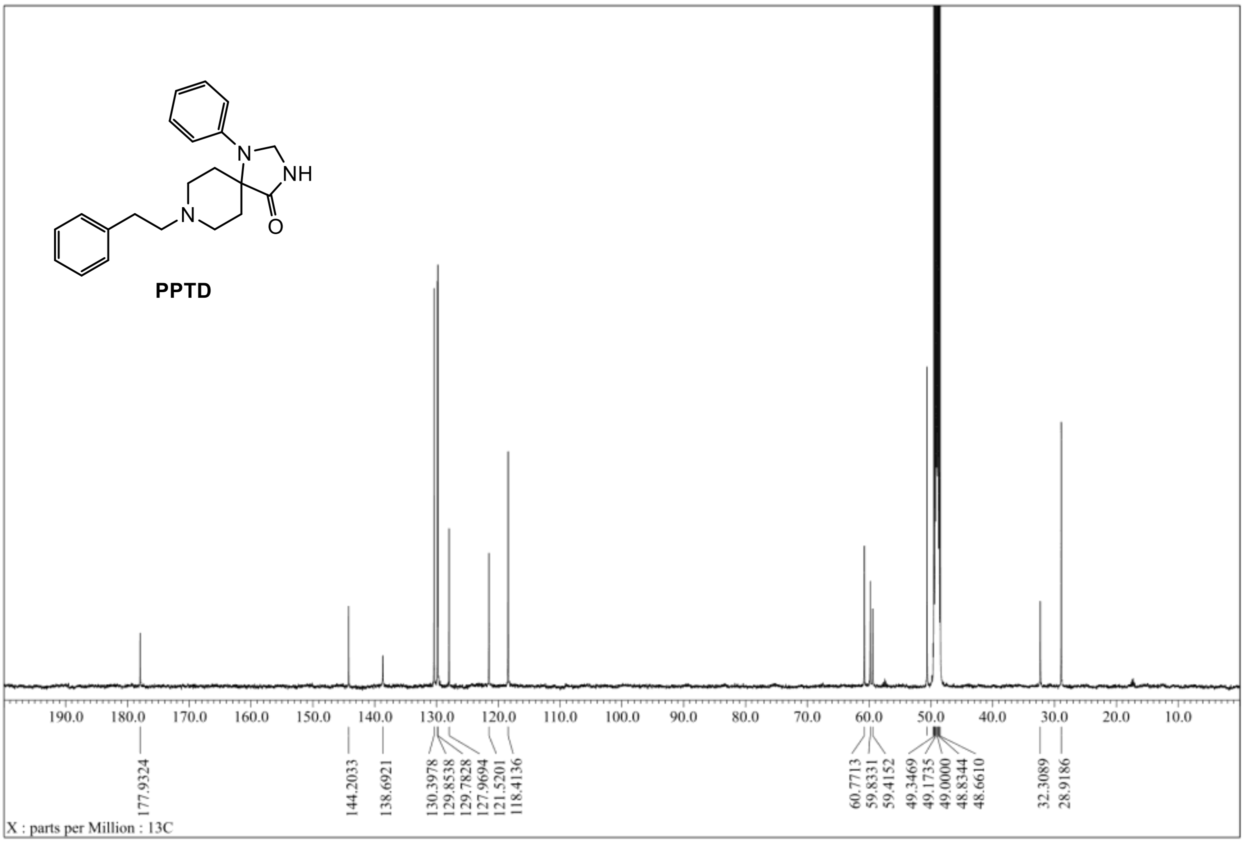


$^{13}$C NMR spectrum of PPTD (125 MHz, $CD_3OD$).

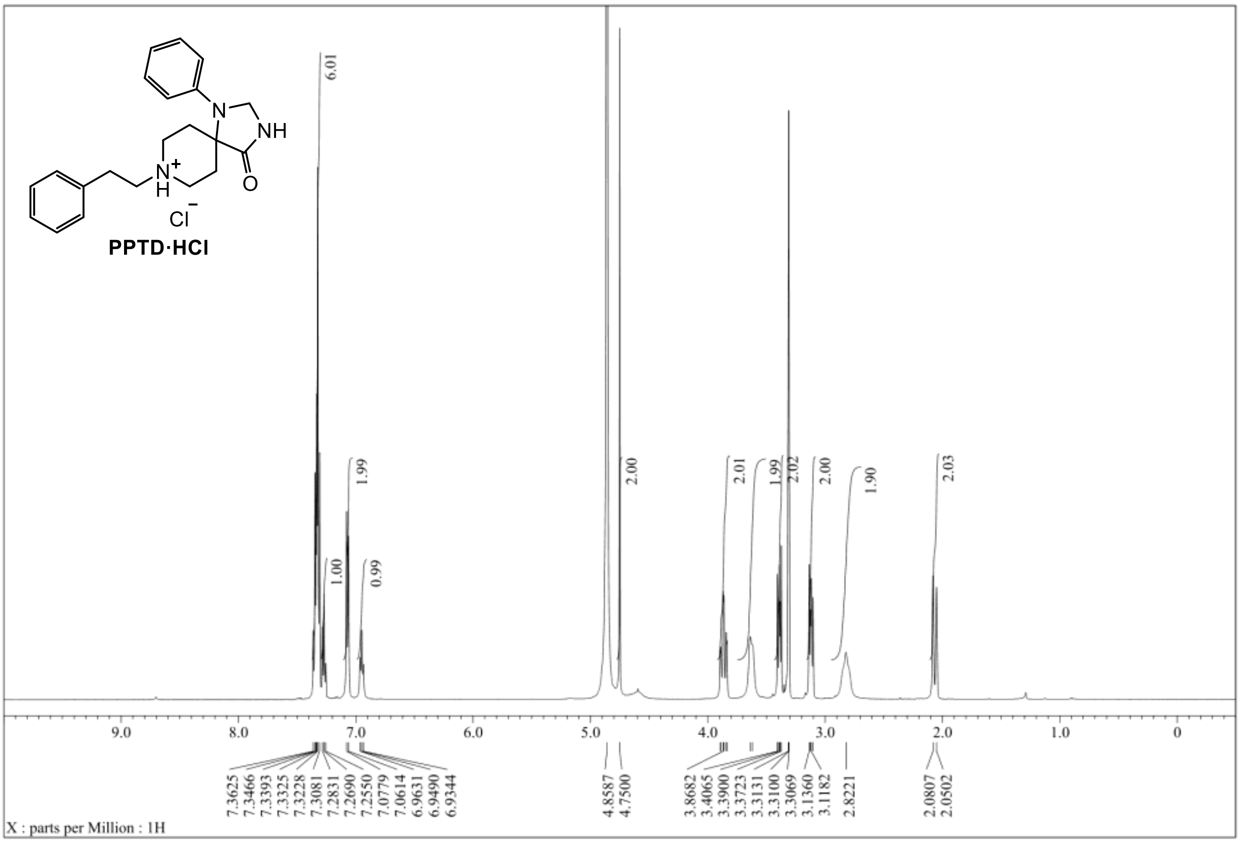


[1]H NMR spectrum of PPTD·HCl (500 MHz, $CD_3OD$).

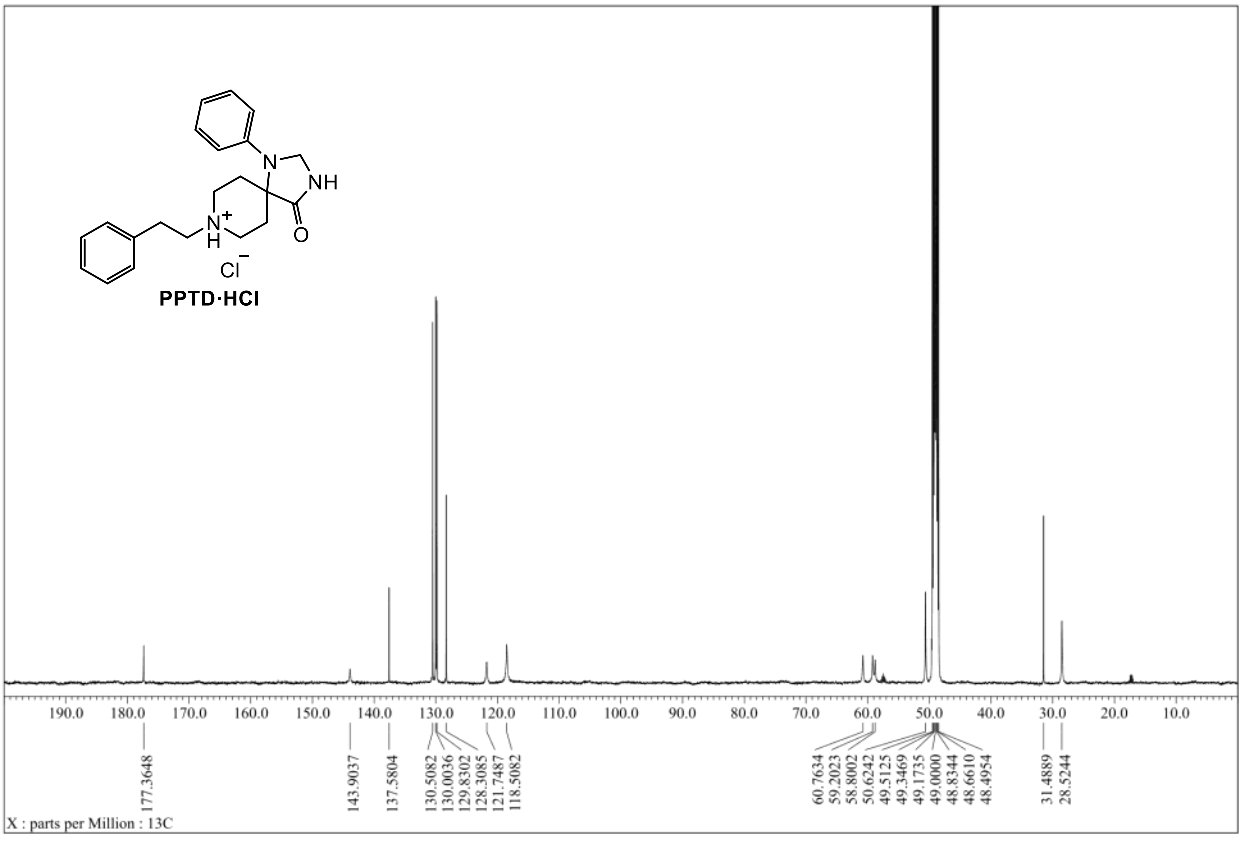


[13]C NMR spectrum of PPTD·HCl (125 MHz, $CD_3OD$).

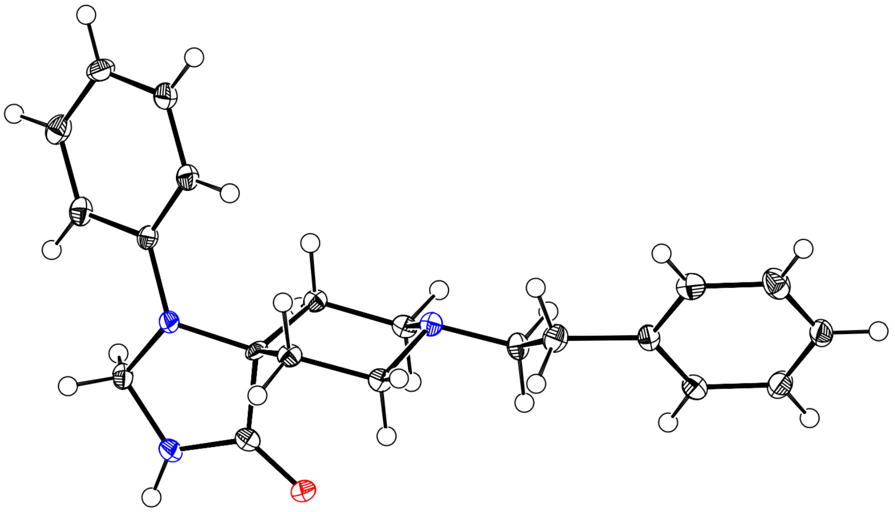

ORTEP drawing of PPTD.

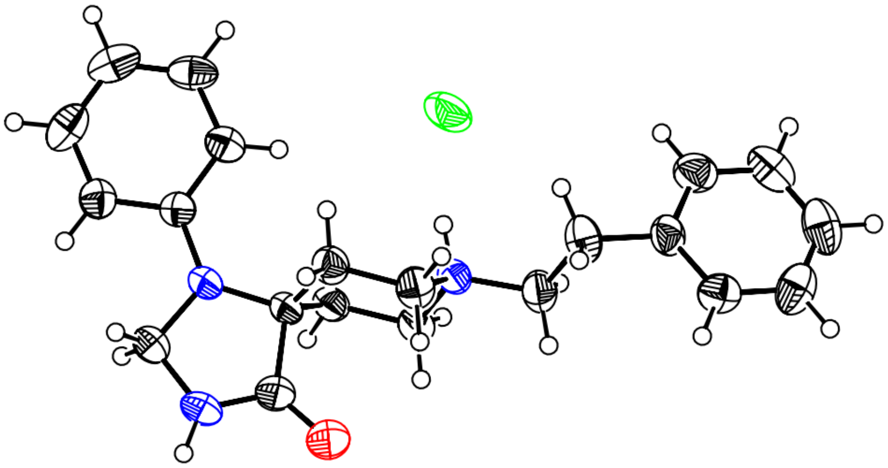

ORTEP drawing of PPTD·HCl.

### Chemical Synthesis of PPTD Derivatives

##### General materials and methods for organic synthesis

All chemical reagents and solvents were obtained from commercial suppliers (Sigma-Aldrich, Tokyo Chemical Industry, FUJIFILM Wako Pure Chemical Corporation, Sasaki Chemical, Nacalai Tesque) and used without further purification. Thin layer chromatography (TLC) was performed on silica gel 60 F254 precoated aluminum sheets or glass plates (Merck) and visualized by fluorescence quenching or ninhydrin staining. Flash column chromatography on silica gel 60 N or Isolera Spektra One (Biotage) was used for chromatographic purification (neutral, 40–50 μm, Kanto Chemical). $^1$H NMR spectra were recorded in deuterated solvents on a JNM-ECS400 (JEOL, 400 MHz) and calibrated to the residual solvent peak or tetramethylsilane (= 0 ppm). Multiplicities are abbreviated as follows: s = singlet, brs = broad singlet, d = doublet, t = triplet, q = quartet, m = multiplet, dd = double doublet. High-resolution mass spectra were measured on an Exactive (Thermo Scientific) equipped with electron spray ionization (ESI). Silicagel 70 F254 PLC Plate was used for preparative TLC purification (PLC) (1 mm thickness, FUJIFILM Wako Pure Chemical Corporation).

##### Synthesis of N1-1 – N1-3

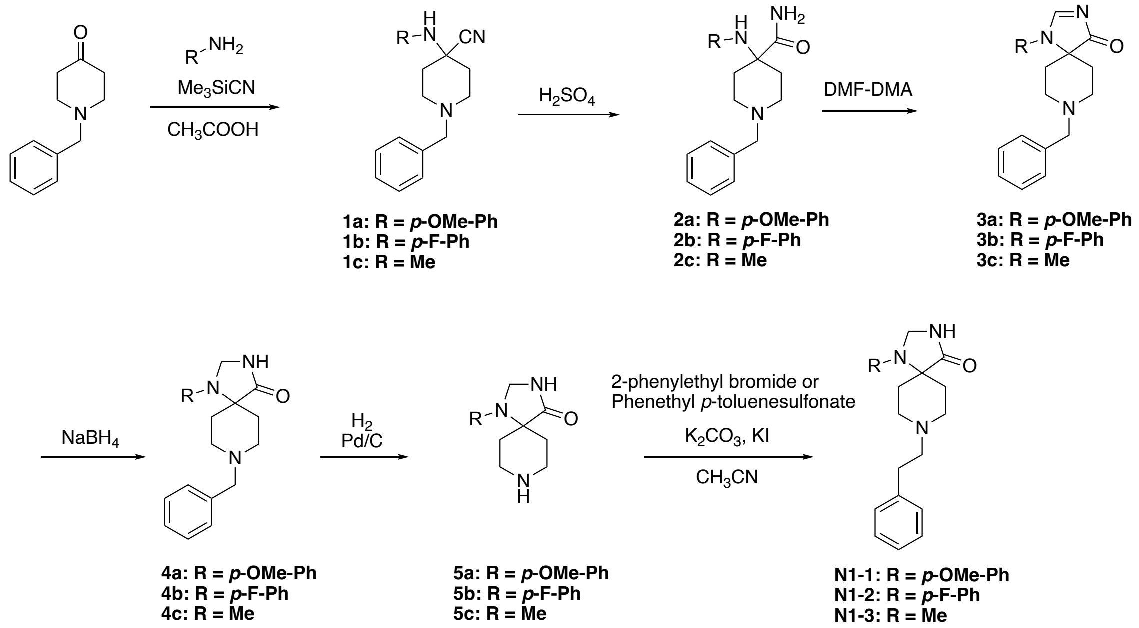


### *1a*

To a stirred solution of 1-benzylpiperidin-4-one (4.50 g, 23.8 mmol) in AcOH (25 mL), 4-methoxyaniline (2.45 g, 19.9 mmol) and $Me_3SiCN$ (3.0 mL, 24.2 mmol) were added at 0 °C. The reaction mixture was stirred at room temperature for 3.5 h and quenched with 2N NaOH (225 mL). The solution was extracted with $CH_2Cl_2$ (50 mL x2), and the organic layer was washed with brine (50 mL x2), dried over $MgSO_4$, filtered, and evaporated.

Diethyl ether (50 mL) was added to the residue, and the resultant solid was filtered and dried in vacuo, resulting in a grey powder of **Int-4** (4.39 g, 68.7%).

$^{1}$**H NMR** (400 MHz, $CDCl_3$): δ/ppm 7.33–7.30 (m, 3H), 7.26–7.24 (m, 2H), 6.98 (d, *J* = 8.8 Hz, 2H), 6.83 (d, *J* = 8.8 Hz, 2H), 3.77 (s, 3H), 3.55 (s, 2H), 3.30 (s, 1H), 2.86–2.83 (m, 2H), 2.43–2.37 (m, 2H), 2.18–2.14 (m, 2H), 1.94–1.87 (m, 2H).

**ESI-MS**: cald. for $[M+H]^+$ as 322.1914, obsd. 322.1915.

***2a***

**1a** (4.332 g, 13.45 mmol) was added to $H_2SO_4$ (23 mL) at room temperature, and stirred for 22 h. The reaction was quenched by adding 28% $NH_3$ aq (75 mL) on ice. The solution was extracted with $CHCl_3$ (75 mL × 2), and the organic layer was washed with brine, dried over $Na_2SO_4$, filtered, and evaporated. Diisopropyl ether (25 mL) was added to the residue, and the resultant solid was filtered and dried in vacuo, resulting in **2a** as a white powder (3.951 g, 86.5%).

$^{1}$**H NMR** (400 MHz, $CDCl_3$): δ/ppm 7.30–7.21 (m, 5H), 7.02 (s, 1H), 6.87 (d, *J* = 8.8 Hz, 2H), 6.59 (d, *J* = 8.8 Hz, 2H), 5.33 (s, 2H), 3.77 (s, 3H), 3.48 (s, 2H), 2.73 (d, *J* = 12.0 Hz, 2H), 2.31 (t, *J* = 12.0 Hz, 2H), 2.08 (t, *J* = 11.2 Hz, 2H), 1.87 (d, *J* = 12.0 Hz, 2H).

**ESI-MS**: cald. for $[M+Na]^+$ as 362.1839, obsd. 362.1845.

***3a***

To a stirred solution of **2a** (3.842 g, 11.32 mmol) in MeOH (22 mL), DMF-DMA (5.0 mL) was added under an atmosphere of $N_2$. The reaction mixture was stirred at 55 °C for 17 h. The solvent was removed in vacuo, and diisopropyl ether (20 mL) was added to the residue. The resultant solid was filtered and washed with diisopropyl ether and diethyl ether, resulting in a grey powder of **3a** (4.083 g, quant).

$^{1}$**H NMR** (400 MHz, $CDCl_3$): δ/ppm 8.16 (s, 1H), 7.30–7.26 (m, 5H), 7.09 (d, *J* = 8.8 Hz, 2H), 6.96 (d, *J* = 8.8 Hz, 2H), 3.85 (s, 3H), 3.57 (s, 2H), 3.03–2.96 (m, 2H), 2.68–2.62 (m, 2H), 1.98–1.91 (m, 2H), 1.79–1.69 (m, 2H).

**ESI-MS**: cald. for $[M+Na]^+$ as 372.1682, obsd. 372.1686.

***4a***

To a stirred solution of **3a** (3.93 g, 11.25 mmol) in dry MeOH (160 mL), $NaBH_4$ (553 mg, 14.6 mmol) was added under an atmosphere of $N_2$. The reaction mixture was stirred at room temperature for 21 h and quenched by adding water (5 mL). The solvent was removed in vacuo, and water (100 mL) was added to the residue. The resultant precipitate was filtered and dried in vacuo at 55 °C, resulting in a grey powder of **4a** (3.241 g, 79.7%).

$^{1}$**H NMR** (400 MHz, DMSO-*d*6): δ/ppm 8.48 (s, 1H), 7.32–7.20 (m, 5H), 6.97 (d, *J* = 8.8 Hz, 2H), 6.89 (d, *J* = 8.8 Hz, 2H), 4.49 (s, 2H), 3.71 (s, 3H), 3.45 (s, 2H), 2.69–2.57 (m, 4H), 2.02–1.94 (m, 2H), 1.64–1.60 (m, 2H).

**ESI-MS**: cald. for $[M+Na]^+$ as 374.1839, obsd. 374.1838.

***5a***

To a stirred solution of **4a** (3.00 g, 8.536 mmol) in MeOH (265 mL) and AcOH (2.5 mL), 10% Pd/C (600 mg) was added. The reaction mixture was stirred under an atmosphere of $H_2$ at room temperature for 24 h. After filtration, the filtrate was evaporated and dissolved in sat $NaHCO_3$ + NaCl aq (35 mL). The solution was extracted with n-BuOH (40 mL × 3), and the organic layer was evaporated. Diethyl ether (50 mL) was added to the residue, and the resultant precipitate was filtered and dried in vacuo at 50 °C, resulting in a white powder of **5a** (2.21 g, 99.1%).

$^1$**H NMR** (400 MHz, DMSO-*d6*): δ/ppm 8.61 (s, 1H), 6.98 (d, *J* = 8.8 Hz, 2H), 6.86 (d, *J* = 8.8 Hz, 2H), 4.52 (s, 2H), 3.70 (s, 3H), 3.25–3.19 (m, 2H), 2.93–2.91 (m, 2H), 2.09–2.02 (m, 2H), 1.65–1.63 (m, 2H).

**ESI-MS**: cald. for $[M+H]^+$ as 262.1550, obsd. 262.1547.

***N1-1***

To a solution of **5a** (0.2038 g, 0.765 mmol) in super dehydrated MeCN (10 mL), dry crushed $K_2CO_3$ (0.3180g, 2.304 mmol), 2-phenylethyl bromide (0.1416 g, 0.765 mmol), and potassium iodine (0.0399 g, 0.240 mmol) were added. The mixture was refluxed for 2 h. The mixture was filtered, and the filtrate was collected and concentrated using a rotary evaporator. The crude was purified on Isolera (Sfar HC Duo 25 g, $CHCl_3$&MeOH: 0 % →5%→10%), and the target-containing fractions were collected, concentrated, and dried in vacuo, yielding a white solid of **N1-1** (270.8 mg, 95 %).

$^1$**H NMR** (400 MHz, $CD_3OD$): δ/ppm 7.28–7.24 (m, 2H), 7.20–7.18 (m, 3H), 7.13 (d, *J* = 8.8 Hz, 2H), 6.89 (d, *J* = 8.8 Hz, 2H), 4.63 (s, 2H), 3.76 (s, 3H), 3.06–2.97 (m, 4H), 2.84–2.72 (m, 4H), 2.08–2.00 (m, 2H), 1.90 (d, *J* = 14.5 Hz, 2H).

**ESI-MS**: cald. for $[M+H]^+$ as 366.2176, obsd. 366.2180.

***1b***

To a stirred solution of 1-benzylpiperidin-4-one (4.50 g, 23.8 mmol) in AcOH (25 mL), 4-fluoroaniline (2.22 g, 19.9 mmol) and $Me_3SiCN$ (3.0 mL, 24.2 mmol) were added at 0 °C. The reaction mixture was stirred at room temperature for 3.5 h and quenched with 2N NaOH (225 mL). The solution was extracted with $CH_2Cl_2$ (50 mL x3), and the organic layer was washed with brine (50 mL x2), dried over $Na_2SO_4$, filtered, and evaporated. Diethyl ether (50 mL) was added to the residue, and the resultant solid was filtered and dried in vacuo, resulting in a pale-yellow powder of **1b** (5.31 g, 86%).

$^1$**H NMR** (400 MHz, $CDCl_3$): δ/ppm 7.35–7.24 (m, 5H), 6.99–6.91 (m, 4H), 3.56 (s, 2H), 2.85–2.81 (m, 2H), 2.46–2.39 (m, 2H), 2.25–2.21 (m, 2H), 1.94–1.87 (m, 2H).

**ESI-MS**: cald. for $[M+H]^+$ as 310.1714, obsd. 310.1715.

***2b***

**1b** (5.26 g, 17.0 mmol) was added to $H_2SO_4$ (27 mL) at room temperature, and the mixture was stirred for 22 h. The reaction was quenched by adding 28% $NH_3$ aq (100 mL) on ice. The solution was extracted with $CHCl_3$ (75 mL x2), and the organic layer was washed with brine, dried over $Na_2SO_4$, filtered, and evaporated. Diisopropyl ether (25 mL) was added to the residue, and the resultant solid was filtered and dried in vacuo, yielding a white powder of **2b** (4.88 g, 88%).

**$^1$H NMR** (400 MHz, $CDCl_3$): δ/ppm 7.31–7.22 (m, 5H), 6.91–6.86 (m, 2H), 6.60–6.55 (m, 2H), 5.34 (s, 1H), 3.93 (s, 1H), 3.49 (s, 2H), 2.76–2.72 (m, 2H), 2.36–2.28 (m, 2H), 2.11–2.04 (m, 2H), 1.89–1.86 (m, 2H).

**ESI-MS**: cald. for $[M+Na]^+$ as 350.1639, obsd. 350.1645.

***3b***

To a stirred solution of **2b** (4.803 g, 14.67 mmol) in MeOH (27 mL), DMF-DMA (6.5 mL) was added under an atmosphere of $N_2$. The reaction mixture was stirred at 55 °C for 17 h. The solvent was removed in vacuo, and diisopropyl ether (20 mL) was added to the residue. The resultant solid was filtered and washed with diisopropyl ether and diethyl ether, yielding **3b** as a pale-yellow powder (4.54 g, 92%).

**$^1$H NMR** (400 MHz, $CDCl_3$): δ/ppm 8.20 (s, 1H), 7.31–7.21 (m, 5H), 7.19–7.16 (m, 4H), 3.57 (s, 2H), 3.04–2.97 (m, 2H), 2.69–2.66 (m, 2H), 1.98–1.91 (m, 2H), 1.81–1.77 (m, 2H).

**ESI-MS**: cald. for $[M+Na]^+$ as 360.1483, obsd. 360.1487.

***4b***

To a stirred solution of **3b** (4.33 g, 12.8 mmol) in dry MeOH (140 mL), $NaBH_4$ (630 mg, 16.7 mmol) was added in an atmosphere of $N_2$. The reaction mixture was stirred at room temperature for 22 h and quenched by adding water (5 mL). The solvent was removed in vacuo, and water (100 mL) was added to the residue. The resultant precipitate was filtered and dried in vacuo at 55 °C, resulting in a white powder of **4b** (3.70 g, 85%).

**$^1$H NMR** (400 MHz, $CDCl_3$): δ/ppm 7.35–7.22 (m, 5H), 7.04–6.93 (m, 4H), 4.67 (s, 2H), 3.56 (s, 2H), 2.82–2.75 (m, 4H), 2.35–2.27 (m, 2H), 1.77–1.73 (m, 2H).

**ESI-MS**: cald. for $[M+Na]^+$ as 362.1639, obsd. 362.1641.

***5b***

To a stirred solution of **4b** (2.00 g, 5.89 mmol) in MeOH (180 mL) and AcOH (2.0 mL), 10% Pd/C (400 mg) was added. The reaction mixture was stirred under an atmosphere containing $H_2$ at room temperature for 24 h. After filtration, the filtrate was evaporated and dissolved in sat $NaHCO_3$ + NaCl aq (35 mL). The solution was extracted with n-BuOH (40 mL x3), and the organic layer was evaporated, yielding **5b** as a pale-yellow powder (1.42 g, 97%).

$^1$**H NMR** (400 MHz, DMSO-*d6*): δ/ppm 8.57 (s, 1H), 7.07 (dd, *J* = 7.2, 8.4 Hz, 2H), 6.92 (dd, *J* = 7.2, 3.6 Hz, 2H), 4.54 (s, 2H), 3.14–3.09 (m, 2H), 2.83–2.80 (m, 2H), 2.23–2.17 (m, 2H), 1.51–1.48 (m, 2H).
**ESI-MS**: cald. for $[M+H]^+$ as 250.1350, obsd. 250.1351.

***N1-2***

To a solution of **5b** (60 mg, 0.24 mmol) in toluene (10 mL), phenethyl *p*-toluenesulfonate (81.9 mg, 0.296 mmol) was added. The reaction mixture was stirred at 80°C for 72 h. After cooling to room temperature, the mixture was filtered and the filtrate was concentrated under reduced pressure. The crude product was purified by flash chromatography (Isolera, Sfar HC Duo 25 g, $CHCl_3$/MeOH gradient: 0% → 5% → 10%), and the target-containing fractions were collected and concentrated in vacuo, yielding **N1-2** as a white solid (23.5 mg, 27%).

$^1$**H-NMR** (400 MHz, $CDCl_3$): δ/ppm 7.31–7.18 (m, 5H), 7.03–6.92 (m, 5H), 6.19 (s, 1H), 4.68 (s, 2H), 2.88–2.79 (m, 6H), 2.75–2.66 (m, 2H), 2.37–2.29 (m, 2H), 1.81–1.78 (m, 2H).
**ESI-MS**: cald. for $[M+H]^+$ as 354.1976, obsd. 354.1972.

***1c***

To a stirred solution of 1-benzylpiperidin-4-one (9.00 g, 47.5 mmol) in AcOH (50 mL), methylamine hydrochloride (4.72 g, 69.9 mmol) and $Me_3SiCN$ (6.0 mL, 48.6 mmol) were added at 0°C. The reaction mixture was stirred at room temperature for 22 h and quenched with 2N NaOH (470 mL). The solution was extracted with $CH_2Cl_2$ (150 mL × 3), and the organic layer was washed with brine (50 mL × 2), dried over $Na_2SO_4$, filtered, and evaporated. Diisopropyl ether (32 mL) and pentane (32 mL) were added to the residue, and the mixture was incubated at -20 °C. The resultant solid was filtered and dried in vacuo, yielding **1c** as a crude product (4.18 g), which was used subsequently without purification.

***2c***

1c (crude, 4.18 g) was added to $H_2SO_4$ (23 mL) at room temperature, and the mixture was stirred for 18 h. The reaction was quenched by adding 28% $NH_3$ aq (65 mL) on ice. The solution was extracted with $CHCl_3$ (50 mL × 3), and the organic layer was washed with brine, dried over $Na_2SO_4$, filtered, and evaporated. Diisopropyl ether (100 mL) was added to the residue, and the resultant solid was filtered and dried in vacuo, yielding **2c** as a white powder (2.05 g, 17% over two steps).
$^1$**H NMR** (400 MHz, $CDCl_3$): δ/ppm 7.33–7.23 (m, 5H), 7.11 (s, 1H), 3.50 (s, 2H), 2.73–2.67 (m, 2H), 2.27 (s, 3H), 2.16–2.11 (m, 4H), 1.67–1.59 (m, 2H).
**ESI-MS**: cald. for $[M+H]^+$ as 248.1757, obsd. 248.1756.

***3c***

To a stirred solution of **2c** (2.02 g, 8.17 mmol) in MeOH (15 mL), DMF-DMA (3.6 mL) was added under $N_2$ atmosphere. The reaction mixture was stirred at 55 °C for 16 h. The solvent was removed in vacuo, and diethyl ether (15 mL) was added to the residue. The resultant solid was filtered and washed with diisopropyl ether and diethyl ether, yielding **3c** as a pale-yellow powder (1.95 g, 93%).

**$^1$H NMR** (400 MHz, $CDCl_3$): δ/ppm 8.06 (s, 1H), 7.36–7.26 (m, 5H), 3.63 (s, 2H), 3.11 (s, 3H), 3.03–2.96 (m, 2H), 2.76–2.71 (m, 2H), 2.04–1.97 (m, 2H), 1.63–1.59 (m, 2H).

**ESI-MS**: cald. for $[M+Na]^+$ as 280.1420, obsd. 280.1421.

***4c***

To a stirred solution of **3c** (1.82 g, 7.06 mmol) in dry MeOH (80 mL), $NaBH_4$ (350 mg, 9.25 mmol) was added under $N_2$ atmosphere. The reaction mixture was stirred at room temperature for 18 h and quenched by adding water (5 mL). The solvent was removed in vacuo, and water (5 mL) was added to the residue. The resultant precipitate was filtered and dried in vacuo at 50°C, yielding **4c** as a white powder (787 mg, 43%).

**$^1$H NMR** (400 MHz, $CDCl_3$): δ/ppm 7.35–7.29 (m, 5H), 4.11 (s, 2H), 3.55 (s, 2H), 2.86–2.22 (m, 2H), 2.69–2.63 (m, 2H), 2.38 (s, 3H), 1.85–1.78 (m, 2H), 1.72–1.68 (m, 2H).

**ESI-MS**: cald. for $[M+Na]^+$ as 282.1577, obsd. 282.1576.

***5c***

To a stirred solution of **4c** (735 mg, 2.83 mmol) in MeOH (50 mL), 10% Pd/C (150 mg) was added. The reaction mixture was stirred at room temperature for 22 h in an atmosphere of $H_2$. After filtration, the filtrate was evaporated, yielding **5c** as colorless oil (532 mg, quant).

**$^1$H NMR** (400 MHz, DMSO-*d*6): δ/ppm 8.04 (s, 1H), 3.92 (s, 2H), 3.03–2.97 (m, 2H), 2.74–2.71 (m, 2H), 2.23 (s, 3H), 1.51–1.41 (m, 4H).

**ESI-MS**: cald. for $[M+H]^+$ as 170.1288, obsd. 170.1287.

***N1-3***

To a solution of **5c** (520 mg, 3.07 mmol) in super dehydrated MeCN (10 mL), 2-phenylethyl bromide (1.12 g, 6.07 mmol) was added. The mixture was refluxed for 46 h. The mixture was filtered, and the solid was dissolved in $NaHCO_3$ aq (50 mL). The solution was extracted with $CHCl_3$ (50 mL x3), and the collected organic layer was washed with brine and dried over $Na_2SO_4$. The organic layer was evaporated, yielding **N1-3** (328 mg, 39 %) as a white powder.

**$^1$H NMR** (400 MHz, $CDCl_3$): δ/ppm 7.31–7.28 (m, 2H), 7.23–7.19 (m, 3H), 5.92 (s, 1H), 4.13 (s, 2H), 3.10–2.75 (m, 6H), 2.40 (s, 3H), 1.78–1.53 (m, 4H).

**ESI-MS**: cald. for $[M+H]^+$ as 274.1914, obsd. 274.1916.

#### Synthesis of N3-1 – N3-3

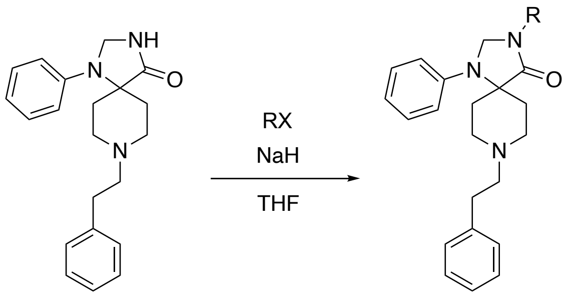


**PPTD**

**N3-1: R = Me**
**N3-2: R = n-propyl**
**N3-3: R = benzyl**

***N3-1***

To a solution of **PPTD** (129 mg, 0.384 mmol) in THF (9 mL), NaH (30.7 mg, 60%, dispersion in Paraffin Liquid, 0.768 mmol) was added at 0 °C in an ice bath. The mixture was stirred at 0 °C for 1 h, and methyl iodide (109 mg, 0.768 mmol) was added. The mixture was stirred at room temperature for 18 h. The mixture was diluted with ethyl acetate (70 mL) and quenched with $H_2O$ (25 mL). The solvent was removed via rotary evaporation, and the crude was purified on Isolera (Sfar HC Duo 25g, $CHCl_3$&MeOH: 0 % →5%→10%). The target-containing fractions were collected, concentrated, and dried in vacuo, yielding **N3-1** (107.7 mg, 80.2 %) as a pale-yellow oil.

**$^1$H-NMR** (400 MHz,$CDCl_3$): δ/ppm 7.30 (m, 4H), 7.24-7.18 (m, 3H), 6.93 (d, $J$ = 8.0 Hz, 2H), 6.87 (t, $J$ = 7.2 Hz, 1H), 4.68 (s, 2H), 3.00-2.76 (m, 13H), 1.70 (d, $J$ = 13.2 Hz, 2H).

**ESI-MS**: cald. for $[M+H]^+$ as 350.2227, obsd.350.2233.

**N3-2**

To a solution of **PPTD** (129 mg, 0.384 mmol) in THF (9 mL), NaH (75.5 mg, 1.89 mmol) was added at 0 °C in an ice bath. The mixture was stirred at 0 °C for 1 h, and 1-iodopropane (261 mg, 1.54 mmol) was added. The mixture was stirred at room temperature for 72 h. The mixture was diluted with ethyl acetate (70 mL) and $H_2O$ (30 mL). The organic layer was collected, and the solvent was removed via rotary evaporation. The crude was purified on Isolera (Sfar HC Duo 25g, $CHCl_3$&MeOH: 0 % →5%→10%). The target-containing fractions were collected, concentrated, and dried in vacuo, yielding **N3-2** (63.5 mg, 44 %) as a pale-yellow oil.

**$^1$H-NMR** (400 MHz, $CD_3OD$) : δ/ppm 7.29–7.15 (m, 7H), 7.04 (d, $J$ = 8.0 Hz, 2H), 6.86 (t, $J$ = 7.4 Hz, 1H), 4.73 (s, 2H), 3.39 (t, $J$ = 7.2 Hz, 2H), 3.00–2.91 (m, 4H), 2.87–2.77 (m, 2H), 2.75–2.60 (m, 4H), 1.81–1.61 (m, 4H), 0.96 (t, $J$ = 7.4 Hz, 3H).

**ESI-MS**: cald. for $[M+H]^+$ as 378.2540, obsd. 378.2540.

**N3-3**

To a solution of **PPTD** (129 mg, 0.384 mmol) in THF (9 mL), NaH (61.6 mg, 1.54 mmol) was added at 0 °C in an ice bath. The mixture was stirred at 0 °C for 1 h, and benzyl bromide (131.4 mg, 0.768 mmol) was added. The mixture was stirred at room temperature for 14 h. The mixture was diluted with ethyl acetate (70 mL) and $H_2O$ (25 mL). The organic layer was washed once with brine (25 mL). The solvent was removed via rotary evaporation. The crude was purified on Isolera (Sfar HC Duo 25g, $CHCl_3$&MeOH: 0 % →5%→10%). The target-containing fractions were collected, concentrated, and dried in vacuo, yielding **N3-3** (66.7 mg, 41 %) as a pale-yellow oil.

**$^1$H-NMR** (400 MHz, $CD_3OD$) : δ/ppm 7.39–7.16 (m, 12H), 6.97 (d, $J$ = 8.0 Hz, 2H), 6.85 (t, $J$ = 7.4 Hz, 1H), 4.61–4.59 (m, 4H), 3.12–2.92 (m, 4H), 2.88–2.84 (m, 2H), 2.72–2.59 (m, 4H), 1.75 (d, $J$ = 14.4 Hz, 2H).

**ESI-MS**: cald. for $[M+H]^+$ as 426.2540, obsd. 426.2538.

#### Synthesis of N8-3 – N8-25

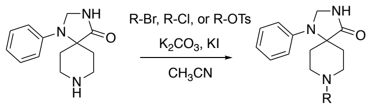


N8-1

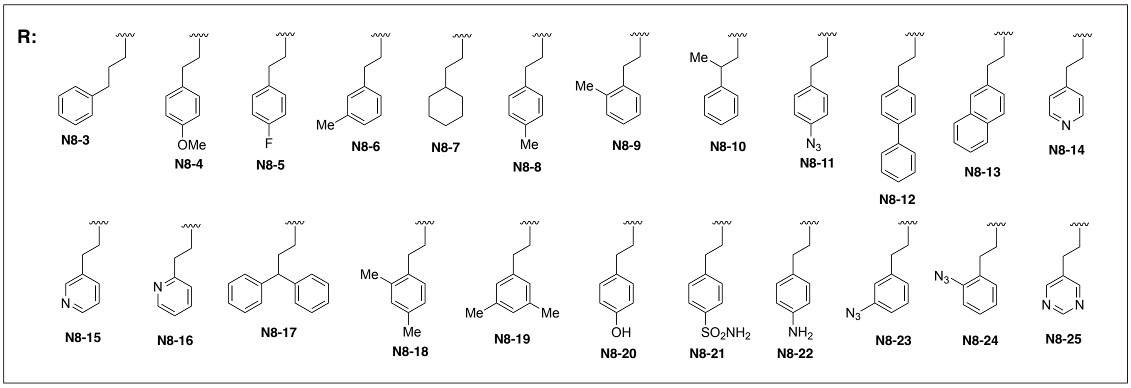


**General protocol for the synthesis of N8 derivatives**

To a solution of commercially available **N8-1** (0.509 g, 2.20 mmol, 1.0 eq) in super dehydrated MeCN (40 mL), dry crushed $K_2CO_3$ (0.898 g, 6.50 mmol, 3.0 eq), alkyl halide or alkyl tosylate (2.20 mmol, 1.0 eq), and potassium iodide (0.107 g, 0.648 mmol, 0.3 eq) were added. The mixture was refluxed for 2 h to overnight. The reaction mixture was filtered, and the filtrate was collected and concentrated using a rotary evaporator. The crude was purified on flash column chromatography (Isolera, Sfar HC Duo 25 g, $CHCl_3$&MeOH: 0 % →5%→10%), and the target-containing fractions were collected, concentrated, and dried in vacuo, yielding the target compound.

***N8-3***

Following the general protocol, 3-phenylpropyl bromide was used as a substrate to obtain **N8-3** (yield: 99.5%).

**$^1$H-NMR** (400 MHz, CDCl3): δ/ppm 7.32–7.24 (m, 4H), 7.24–7.15 (m, 3H), 6.93 (d, *J* = 8.0 Hz, 2H), 6.87 (t, *J* = 7.6 Hz), 6.19 (bs, 1H), 4.73 (s, 2H), 2.84 (bs, 4H), 2.67 (t, *J* = 7.6 Hz, 2H), 2.48 (bs, 2H), 1.88 (bs, 2H), 1.74 (d, *J* = 14 Hz), 1.60 (bs, 2H)

**ESI-MS**: cald. for $[M+H]^+$ as 350.2227, obsd.350.2229.

***N8-4***

Following the general protocol, 4-methoxyphenethyl bromide was used as a substrate to obtain **N8-4** (yield: 79.6%).

**$^1$H-NMR** (400 MHz, $CD_3OD$): δ/ppm 7.27 (td, *J* = 7.0, 2.0 Hz, 2H), 7.14 (dd, *J* = 6.8, 2.0 Hz, 2H), 7.02 (dd, *J* = 8.7, 0.8 Hz, 2H), 6.89–6.84 (m, 3H), 4.70 (s, 2H), 3.76 (s, 3H), 3.16–3.10 (m, 4H), 2.85–2.82 (m, 4H), 2.70–2.62 (m, 2H), 1.82 (d, *J* = 14.4 Hz, 2H).

**ESI-MS**: cald. for $[M+H]^+$ as 366.2176, obsd.366.2180.

***N8-5***

Following the general protocol, 2-(4-fluorophenyl) ethyl bromide was used as a substrate to obtain **N8-5** (yield: 74.1%).

**$^1$H-NMR** (400 MHz, $CD_3OD$): δ/ppm 7.29–7.23 (m, 4H), 7.04–6.99 (m, 4H), 6.88 (t, *J* = 7.2 Hz, 1H), 4.70 (s, 2H), 3.16–3.13 (m, 4H), 2.89–2.83 (m, 4H), 2.69–2.61 (m, 2H), 1.82 (d, *J* = 14.8 Hz, 2H).

**ESI-MS**: cald. for $[M+H]^+$ as 354.1976, obsd.354.1978.

***N8-6***

Following the general protocol, 3-methylphenethyl bromide was used as a substrate to obtain **N8-6** (yield: 80.0%).

**$^1$H-NMR** (400 MHz, $CD_3OD$): δ/ppm 7.29–7.24 (m, 2H), 7.16 (t, *J* = 7.6 Hz, 1H), 7.05–7.00 (m, 5H), 6.87 (t, *J* = 7.2 Hz, 1H), 4.70 (s, 2H), 3.13–3.07 (m, 4H), 2.86–2.80 (m, 4H), 2.70–2.62 (m, 2H), 2.31 (s, 3H), 1.81 (d, *J* = 14.4 Hz, 2H).

**ESI-MS**: cald. for $[M+H]^+$ as 350.2227, obsd. 350.2229.

***N8-7***

Following the general protocol, 2-cyclohexylethyl bromide was used as a substrate to obtain **N8-7** (yield: 30.0%).

**$^1$H-NMR** (400 MHz, $CD_3OD$): δ/ppm 7.30 (t, *J* = 7.8 Hz, 2H), 7.02 (d, *J* = 8.4 Hz, 2H), 6.94 (t, *J* = 7.2 Hz, 1H), 4.72 (s, 2H), 3.62–3.57 (m, 2 H), 3.41–3.38 (m, 2H), 3.05–3.01 (m, 2H), 2.69–2.61 (m, 2H), 1.99 (d, *J* = 14.8 Hz, 2H), 1.76–1.66 (m, 5H), 1.62–1.56 (m, 2H), 1.37–1.18 (m, 4H), 1.05–0.95 (m, 2H).
**ESI-MS**: cald. for $[M+H]^+$ as 342.2540, obsd. 342.2543.

***N8-8***
Following the general protocol, 4-methylphenethyl bromide was used as a substrate to obtain **N8-8** (yield: 75.0%).
**$^1$H-NMR** (400 MHz, DMSO-D6): δ/ppm 8.63 (s, 1H), 7.23 (t, *J* = 8.0 Hz, 2H), 7.13 (d, *J* = 8.4 Hz, 2H), 7.09 (d, *J* = 8.0 Hz, 2H), 6.82 (d, *J* = 8.0 Hz, 2H), 6.75 (t, *J* = 7.2 Hz, 1H), 4.57 (s, 2H), 2.86–2.78 (m, 2H), 2.76–2.67 (m, 4H), 2.59–2.53 (m, 4H), 2.27 (s, 3H), 1.57 (d, *J* = 13.2 Hz, 2H)
**ESI-MS**: cald. for $[M+H]^+$ as 350.2227, obsd. 350.2234.

***N8-9***
Following the general protocol, 2-methylphenethyl bromide was used as a substrate to obtain **N8-9** (yield: 74.8%).
**$^1$H-NMR** (400 MHz, DMSO-D6): δ/ppm 8.64 (s, 1H), 7.23 (t, *J* = 8.0 Hz, 2H), 7.19–7.06 (m, 4H), 6.84 (d, *J* = 8.0 Hz, 2H), 6.75 (t, *J* = 7.2 Hz, 1H), 4.58 (s, 2H), 2.88–2.72 (m, 6H), 2.62–2.52 (m, 4H), 2.30 (s, 3H), 1.58 (d, *J* = 13.2 Hz, 2H)
**ESI-MS**: cald. for $[M+H]^+$ as 350.2227, obsd. 350.2236.

***N8-10***
Following the general protocol, 1-bromo-2-phenylpropane was used as a substrate to obtain **N8-10** (yield: 19.6%).
**$^1$H-NMR** (400 MHz, CDCl3): δ/ppm 7.34–7.18 (m, 7H), 6.87–6.82 (m, 3H), 6.19 (bs, 1H), 4.71 (s, 2H), 3.02–2.92 (m, 1H), 2.84–2.49 (m, 8H), 1.66 (bd, *J* = 16.0 Hz, 2H), 1.32 (d, *J* = 6.8 Hz, 3H)
**ESI-MS**: cald. for $[M+H]^+$ as 350.2227, obsd. 350.2234.

***N8-11***
Following the general protocol, 4-azidophenethyl 4-methylbenzenesulfonate was used as a substrate to obtain **N8-11** (yield: 44.0%).
**$^1$H-NMR** (400 MHz, CDCl3): δ/ppm 7.30 (t, *J* = 8.0 Hz, 2H), 7.21 (d, *J* = 8.4 Hz, 2H), 6.96 (dd, *J* = 6.6Hz, *J* = 2.0 Hz, 2H), 6.96–6.86 (m, 3H), 6.12 (bs, 1H), 4.74 (s, 2H), 2.92–2.86 (m, 4H), 2.84–2.78 (m, 2H), 2.72–2.61 (m, 4H), 1.75 (d, *J* = 14.0 Hz, 2H)

**ESI-MS**: cald. for $[M+H]^+$ as 377.2084, obsd. 377.2087.

***N8-12***

Following the general protocol, 4-(2-bromoethyl)-1,1'-biphenyl was used as a substrate to obtain **N8-12** (yield: 89.9%).

**$^1$H-NMR** (400 MHz, CDCl3): δ/ppm 7.59 (dd, *J* = 8.4 Hz, *J* = 1.2 Hz, 2H), 7.53 (dd, *J* = 6.4 Hz, *J* = 2.0 Hz, 2H), 7.43 (t, *J* = 7.6 Hz, 2H), 7.36–7.26 (m, 5H), 6.94 (d, *J* = 8.0 Hz, 2H), 6.88 (t, *J* = 7.2 Hz, 1H), 6.04 (bs, 1H), 4.74 (s, 2H), 2.96–2.85 (m, 6H), 2.78–2.63 (m, 4H), 1.77 (bd, *J* = 14.0 Hz, 2H)

**ESI-MS**: cald. for $[M+H]^+$ as 412.2383, obsd. 412.2399.

***N8-13***

Following the general protocol, 2-(2-bromoethyl)naphthalene was used as a substrate to obtain **N8-13** (yield: 82.5%).

**$^1$H-NMR** (400 MHz, DMSO-D6): δ/ppm 8.63 (s, 1H), 7.89 (d, *J* = 7.2 Hz, 1H), 7.85 (dd, *J* = 8.0 Hz, *J* = 3.6 Hz, 2H), 7.76 (s, 1H), 7.52–7.42 (m, 3H), 7.18 (t, *J* = 8.2 Hz, 2H), 6.81 (d, *J* = 8.0 Hz, 2H), 6.73 (t, *J* = 7.2 Hz, 1H), 2.98–2.84 (m, 4H), 2.82–2.72 (m, 2H), 2.70–2.63 (m, 2H), 2.61–2.54 (m, 3H), 1.58 (bd, *J* = 13.2 Hz, 2H)

**ESI-MS**: cald. for $[M+H]^+$ as 386.2227, obsd. 386.2230.

***N8-14***

Following the general protocol, 4-(2-bromoethyl)pyridine was used as a substrate to obtain **N8-14** (yield: 28.5%).

**$^1$H-NMR** (400 MHz, CDCl3): δ/ppm 8.51 (dd, *J* = 7.6 Hz, *J* = 1.6 Hz, 2H), 7.29 (t, *J* = 7.6 Hz, 2H), 7.16 (d, *J* = 6.0 Hz, 2H), 6.92–6.85 (m, 3H), 5.98 (bs, 1H), 4.74 (s, 2H), 2.95–2.79 (m, 6H), 2.75–2.61 (m 4H), 1.75 (bd, *J* = 14.0 Hz, 2H)

**ESI-MS**: cald. for $[M+Na]^+$ as 359.1842, obsd. 359.1849.

***N8-15***

Following the general protocol, 3-(2-chloroethyl)pyridine was used as a substrate to obtain **N8-15** (yield: 25.3%).

**$^1$H-NMR** (400 MHz, CDCl3): δ/ppm 8.50 (d, *J* = 2.0 Hz, 1H), 8.47 (dd, *J* = 5.2 Hz, *J* = 1.6 Hz, 1H), 7.55 (dt, *J* = 8.0 Hz, *J* = 2.0 Hz, 1H), 7.29 (t, *J* = 8.0 Hz, 2H), 7.23 (dd, *J* = 7.2 Hz, 4.8 Hz, 1H), 6.94–6.85 (m, 3H), 6.19 (bs, 1H), 4.74 (s, 2H), 2.96–2.86 (m, 4H), 2.86–2.80 (m, 2H), 2.74–2.62(m 4H), 1.75 (bd, *J* = 14.0 Hz, 2H)

**ESI-MS**: cald. for $[M+Na]^+$ as 359.1842, obsd. 359.1844.

***N8-16***

Following the general protocol, 2-(2-bromoethyl)pyridine was used as a substrate to obtain **N8-16** (yield: 70.0%).

**$^1$H-NMR** (400 MHz, CDCl3): δ/ppm 8.54 (dq, *J* = 4.8 Hz, *J* = 1.6 Hz, *J* = 0.8 Hz, 1H), 7.61 (dt, *J* = 7.6 Hz, *J* = 1.6 Hz, 1H), 7.31–7.27 (m, 2H), 7.21 (d, *J* = 7.6 Hz, 1H), 7.13 (dt, *J* = 6.4 Hz, *J* = 1.2 Hz, 1H), 6.92–6.83 (m 3H), 4.74 (s, 2H), 3.06–3.00 (m 2H), 2.96–2.83 (m 6H), 2.72–2.60 (m, 2H), 1.74 (bd, *J* = 14.4 Hz, 2H)

**ESI-MS**: cald. for $[M+H]^+$ as 337.2023, obsd. 337.2021.

***N8-17***

Following the general protocol, 3,3-diphenylpropyl 4-methylbenzenesulfonate was used as a substrate to obtain **N8-17** (yield: 77.4%).

**$^1$H-NMR** (400 MHz, CDCl3): δ/ppm 7.35–7.24 (m, 10H), 7.21–7.14 (m, 2H), 6.93 (d, *J* = 8.0 Hz, 2H), 6.88 (t, *J* = 7.2 Hz, 1H), 6.05 (bs, 1H), 4.72 (s, 2H), 4.03 (t, *J* = 7.6 Hz, 1H), 2.84–2.71 (m, 4H), 2.69–2.57 (m, 2H), 2.41–2.35 (m, 2H), 2.32–2.23 (m, 2H), 1.71 (bd, *J* = 13.6 Hz, 2H)

**ESI-MS**: cald. for $[M+H]^+$ as 426.2540, obsd. 426.2550.

***N8-18***

Following the general protocol, 2,4-dimethylphenethyl 4-methylbenzenesulfonate was used as a substrate to obtain **N8-18** (yield: 58.3%).

**$^1$H-NMR** (400 MHz, CDCl3): δ/ppm 7.28 (t, *J* = 8.0 Hz, 2H), 7.05 (d, *J* = 8.0 Hz, 1H), 6.97–6.20 (m, 4H), 6.89 (t, *J* = 7.2 Hz, 1H), 5.95 (bs, 1H), 4.72 (s, 2H), 2.95–2.88 (m, 4H), 2.83–2.76 (m, 2H), 2.72–2.58 (m, 4H), 2.29 (s, 3H), 2.28(s, 3H), 1.75 (d, *J* = 14.4 Hz, 2H)

**ESI-MS**: cald. for $[M+H]^+$ as 364.2383, obsd. 364.2386.

***N8-19***

Following the general protocol, 3,5-dimethylphenethyl 4-methylbenzenesulfonate was used as a substrate to obtain **N8-19** (yield: 78.1%).

**$^1$H-NMR** (400 MHz, CDCl3): δ/ppm 7.28 (t, *J* = 8.0 Hz, 2H), 6.92 (d, *J* = 7.6 Hz, 2H), 6.87 (t, *J* = 7.6 Hz, 1H), 6.84 (s, 3H), 6.24 (bs, 1H), 4.73 (s, 2H), 2.92–2.82 (m, 4H), 2.78–2.62 (m, 6H), 2.28 (s, 6H), 1.75 (bd, *J* = 14.0 Hz, 2H)

**ESI-MS**: cald. for $[M+H]^+$ as 364.2383, obsd. 364.2388.

***N8-20***

Following the general protocol, 4-(2-bromoethyl)phenol was used as a substrate to obtain **N8-20** (yield: 15.6%).

**¹H-NMR** (400 MHz, DMSO-D6): δ/ppm 9.15 (bs, 1H), 8.64 (bs, 1H), 7.23 (t, *J* = 7.8 Hz, 2H), 7.03 (d, *J* = 8.0 Hz, 2H), 6.83 (d, *J* = 8.0 Hz, 2H), 6.75 (t, *J* = 7.2 Hz, 1H), 6.67 (d, *J* = 8.4 Hz, 2H), 4.58 (s, 2H), 2.90–2.40 (m, 10H), 1.57 (bd, *J* = 12.0 Hz, 2H)
**ESI-MS**: cald. for $[M+H]^+$ as 352.2020, obsd. 352.2028.

***N8-21***
Following the general protocol, 4-(2-bromoethyl)benzenesulfonamide was used as a substrate to obtain **N8-21** (yield: 25.9%).
**¹H-NMR** (400 MHz, DMSO-D6): δ /ppm 8.63 (s, 1H), 7.74 (d, *J* = 8.4 Hz, 2H), 7.44 (d, *J* = 8.4 Hz, 2H), 7.30 (bs, 2H), 7.23 (t, *J* = 8.0 Hz, 2H), 6.81 (d, *J* = 8.0 Hz, 2H), 6.74 (t, *J* = 7.2 Hz), 4.58 (s, 2H), 2.89–2.79 (m, 4H), 2.78–2.66 (m, 2H), 2.62–2.53 (m, 4H), 1.57 (bd, *J* = 13.2 Hz, 2H)
**ESI-MS**: cald. for $[M+H]^+$ as 415.1798, obsd. 415.1799.

***N8-22***
Following the general protocol, 4-((*tert*-butoxycarbonyl)amino)phenethyl 4-methylbenzenesulfonate was used as a substrate to obtain Boc-protected **N8-22**. The compound was deprotected with 50% TFA in $CH_2Cl_2$ to obtain **N8-22** (two-step yield: 68.5%).
**¹H-NMR** (400 MHz, CDCl3): δ/ppm 7.29 (t, *J* = 7.6 Hz, 2H), 7.01 (d, *J* = 8.4 Hz, 2H), 6.93 (d, *J* = 8.0 Hz, 2H), 6.88 (t, *J* = 7.6 Hz, 1H), 6.63 (dd, *J* = 8.4 Hz, 2H), 6.08 (bs, 1H), 4.74 (s, 2H), 3.49 (bs, 2H), 2.94–2.82 (m, 4H), 2.76–2.60 (m, 6H), 1.75 (bd, *J* = 14.4 Hz, 2H)
**ESI-MS**: cald. for $[M+H]^+$ as 351.2179, obsd. 351.2187.

***N8-23***
Following the general protocol, 3-azidophenethyl 4-methylbenzenesulfonate was used as a substrate to obtain **N8-23** (yield: 60.9%).
**¹H-NMR** (400 MHz, CDCl3): δ/ppm 7.33–7.24 (m, 3H), 7.01 (d, *J* = 7.2 Hz, 1H), 6.94–6.86 (m, 5H), 6.13 (bs, 1H), 4.74 (s, 2H), 2.96–2.86 (m, 4H), 2.85–2.78 (m, 2H), 2.73–2.62 (m 4H), 1.75 (bd, *J* = 14.0 Hz, 2H)
**ESI-MS**: cald. for $[M+H]^+$ as 377.2084, obsd. 377.2088.

***N8-24***
Following the general protocol, 2-azidophenethyl 4-methylbenzenesulfonate was used as a substrate to obtain **N8-24** (yield: 37.3%).

**$^{1}$H-NMR** (400 MHz, CDCl3): δ/ppm 7.33–7.24 (m, 3H), 7.21 (dd, *J* = 7.6 Hz, 1.4 Hz, 1H), 7.14 (dd, *J* = 8.0 Hz, *J* = 1.2 Hz, 1H), 7.08 (dt, *J* = 7.2 Hz, 1.2 Hz, 1H), 6.92 (d, *J* = 8.0 Hz, 2H), 6.88 (t, *J* = 7.2 Hz, 1H), 6.27 (bs, 1H), 4.74 (s, 2H), 2.94–2.85 (m, 4H), 2.84–2.77 (m, 2H), 2.72–2.60 (m, 4H), 1.75 (bd, *J* = 14.4 Hz, 2H)
**ESI-MS**: cald. for $[M+Na]^+$ as 399.1904, obsd. 399.1911.

***N8-25***
Following the general protocol, 2-(pyrimidin-5-yl)ethyl 4-methylbenzenesulfonate was used as a substrate to obtain **N8-25** (yield: 93.3%).
**$^{1}$H-NMR** (400 MHz, CDCl3): δ/ppm 9.11 (s, 1H), 8.64 (s, 2H), 7.31 (t, *J* = 8.0 Hz, 2H), 6.91–6.85 (m, 3H), 6.34 (bs, 1H), 4.74 (s, 2H), 2.96–2.78 (m, 6H), 2.74–2.62 (m 4H), 1.75 (d, J = 14.4 Hz, 2H)
**ESI-MS**: cald. for $[M+Na]^+$ as 360.1795, obsd. 360.1792.

**Supplementary Figs**

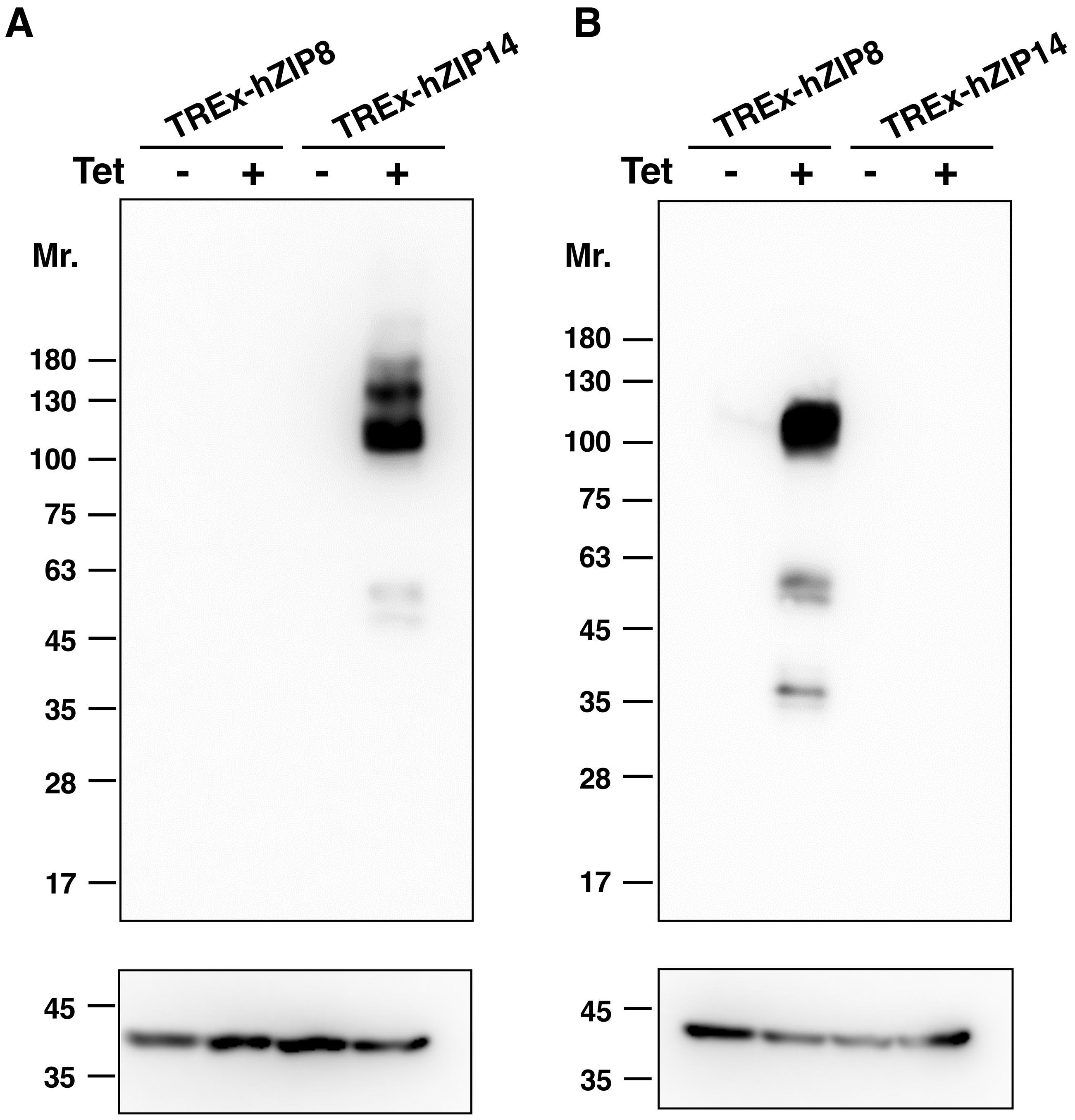


**Supplementary Fig. 1.**

**Expression profiles of ZIP8 and ZIP14 in cells.**

Cell lysates from either TREx-hZIP14 cells or TREx-hZIP8 cells treated with tetracycline (Tet) for 24h were analyzed by western blot, using (**A**) anti-ZIP14, (**B**) anti-ZIP8, or anti-β-actin antibodies as a loading control (lower panels).

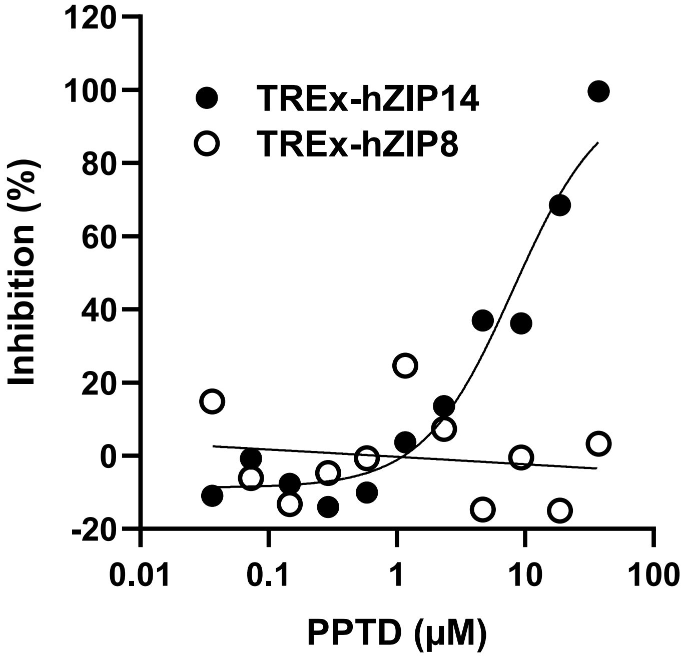


**Supplementary Fig. 2.**

**Effect of PPTD on ZIP14- or ZIP8-mediated zinc influx in TREx-hZIP14 and TREx-hZIP8 cells.**

Intracellular zinc levels were measured using FluoZin-3 AM fluorescence following tetracycline (Tet) induction, as described in the *Methods*. Data are presented as percentages relative to DMSO-treated cells (0%) and $ZnSO_4$-untreated cells (100%). A dose-dependent inhibition of zinc influx by PPTD was observed in ZIP14-expressing cells (black circles), but not in ZIP8-expressing cells (white circles). PPTD exhibited an $IC_{50}$ of 8.1 μM for ZIP14-mediated zinc uptake in TREx-hZIP14 cells, whereas no significant inhibition was observed for ZIP8-mediated uptake in TREx-hZIP8 cells up to 37 μM.

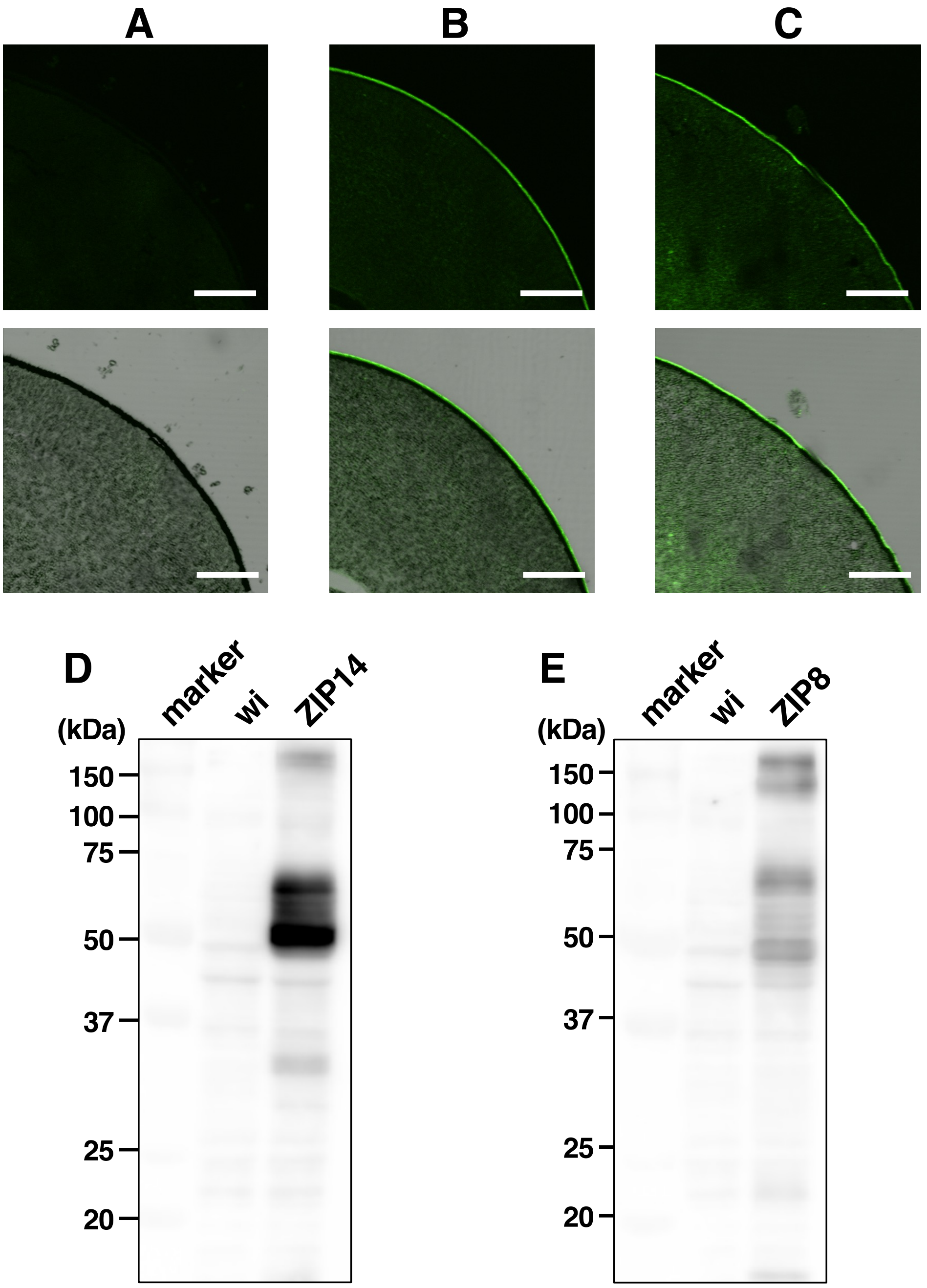


**Supplementary Fig. 3.**

**Expression profiles of ZIP8 and ZIP14 in *Xenopus laevis* oocytes.**

Oocytes were injected with (**A**) either water alone, cRNA encoding (**B**) human ZIP14, or (**C**) human ZIP8, and analyzed by immunofluorescence microscopy using an anti-HA antibody followed by an Alexa Fluor 488-conjugated anti-mouse IgG secondary antibody. Upper panels show immunofluorescence images; lower panels display merged immunofluorescence and differential interference contrast (DIC) images. Total lysates from oocytes injected with water (wi), cRNA encoding (**D**) human ZIP14, or human (**E**) ZIP8 were subjected to western blot analysis using an anti-HA antibody and an HRP-conjugated anti-mouse IgG secondary antibody. *Marker* indicates the molecular weight marker used in the western blot. Scalebar:100µm

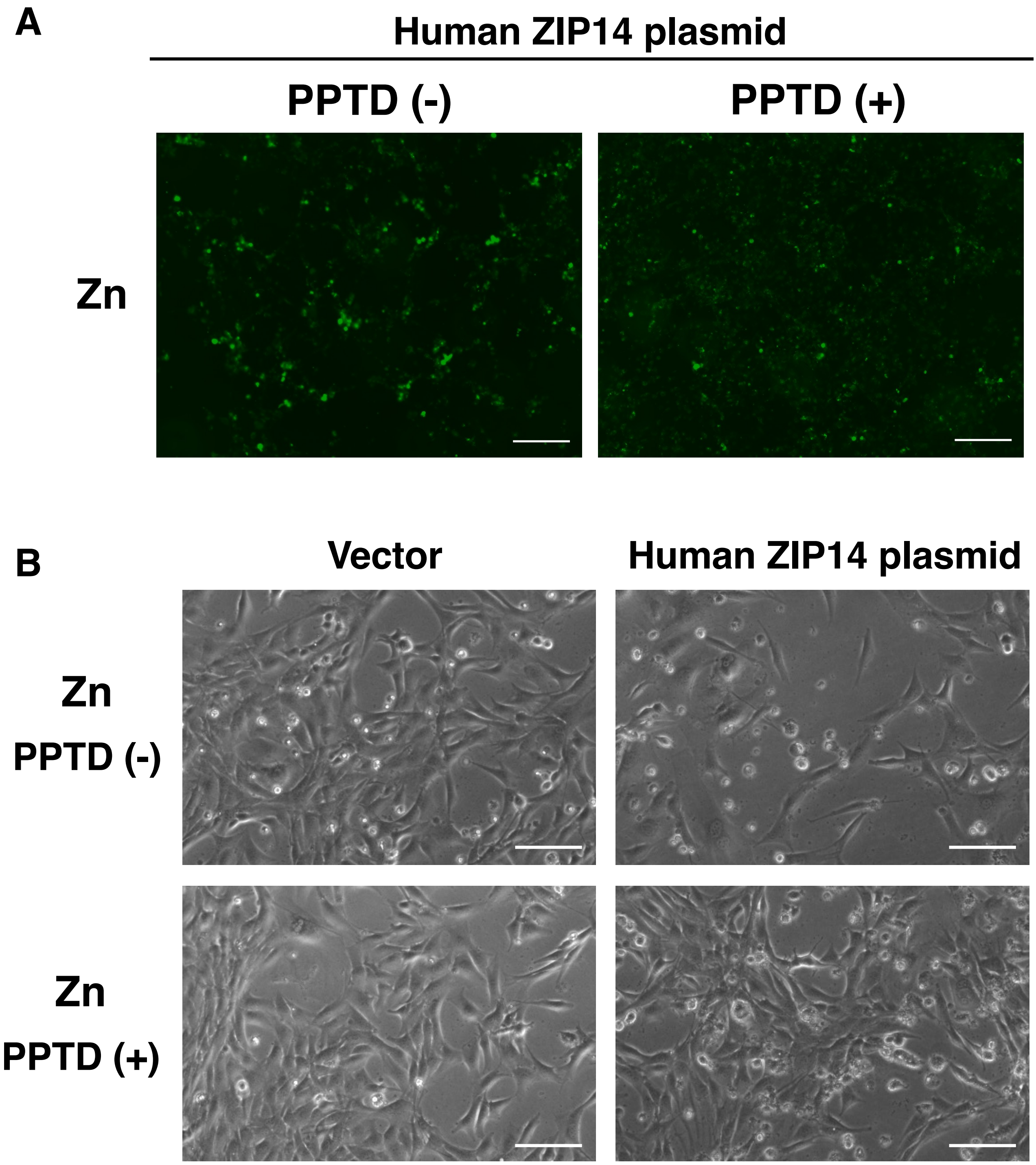


**Supplementary Fig. 4.**

**Effect of PPTD on preventing ZIP14-mediated cytotoxicity in mouse myoblast C2C12 cells.**

ZIP14-mediated zinc influx (**A**, scale: 200 μm) and cytotoxicity (**B**, scale: 30 μm) in C2C12 myoblast cells, which was prevented by PPTD treatment. Human ZIP14 expression plasmid was transfected into C2C12 cells. $ZnSO_4$ (final concentration 100 μM) and PPTD (final concentration 10 μM) were added 24 h post-transfection. Cells were incubated for an additional 5 days at 37°C in 5% $CO_2$. Intracellular zinc level was visualized by Zinpyr-1 (final concentration 0.5 μM).

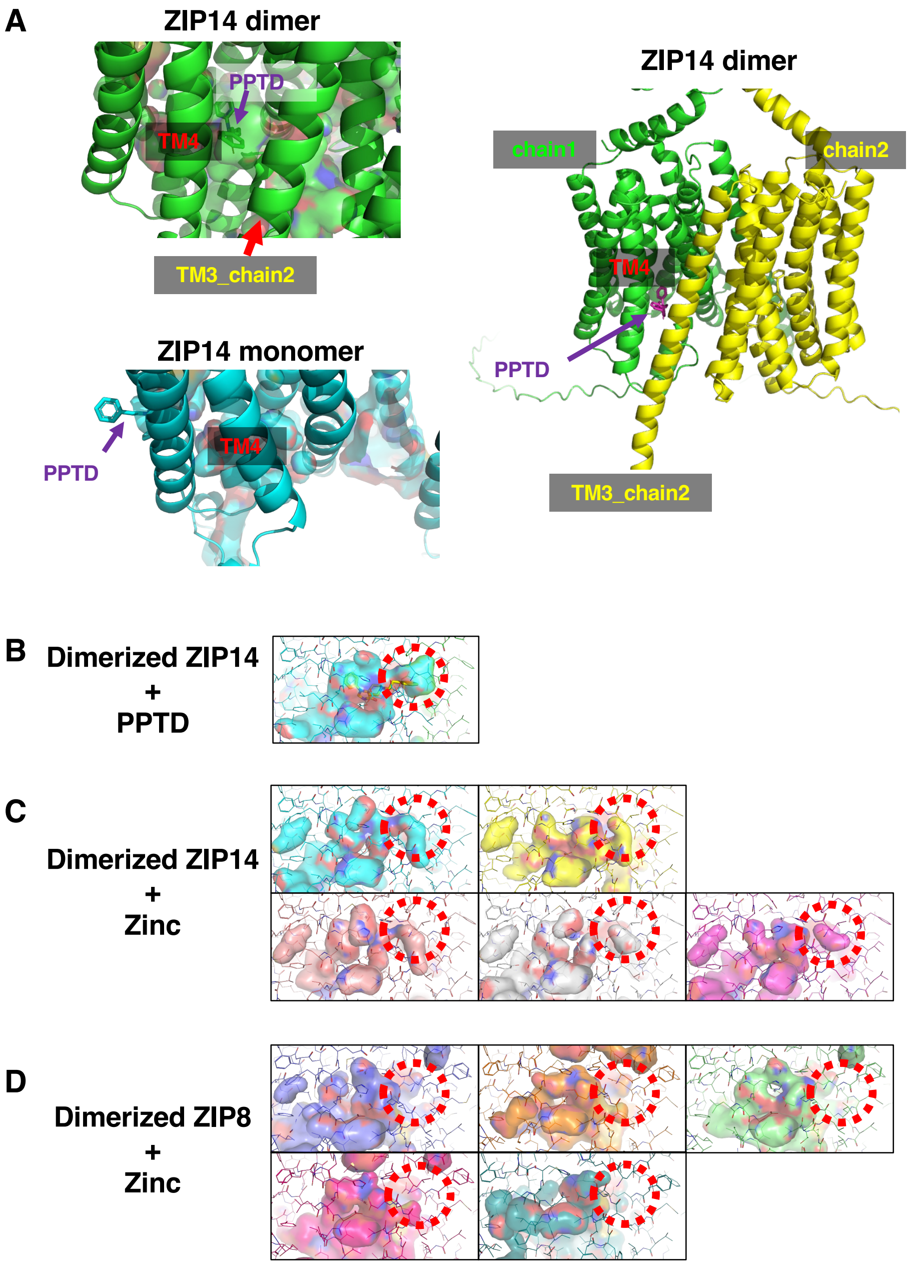


**Supplementary Fig. 5.**

**Predicted binding pocket of PPTD in ZIP14.**

**A** Left: Predicted binding mode of PPTD in dimeric human ZIP14 (left upper) versus monomeric ZIP14 (left lower). Right: A structural rearrangement is observed upon dimerization, in which transmembrane helix 3 (TM3) from one protomer (TM3_chain2) interacts with TM4 of the adjacent protomer, generating an internal pocket that accommodates PPTD (left upper).

**B** Structural model of dimeric human ZIP14 bound to PPTD. A predicted binding pocket for PPTD is indicated by the red dashed circle.

**C** Structural models of five human ZIP14 dimers in complex with zinc. The red dashed circle marks the region corresponding to the PPTD-binding pocket in each model.

**D** Structural models of five human ZIP8 dimers in complex with zinc, showing no comparable pocket formation. Red dashed circles indicate the regions analogous to the PPTD-binding pocket in ZIP14, used for comparative reference.

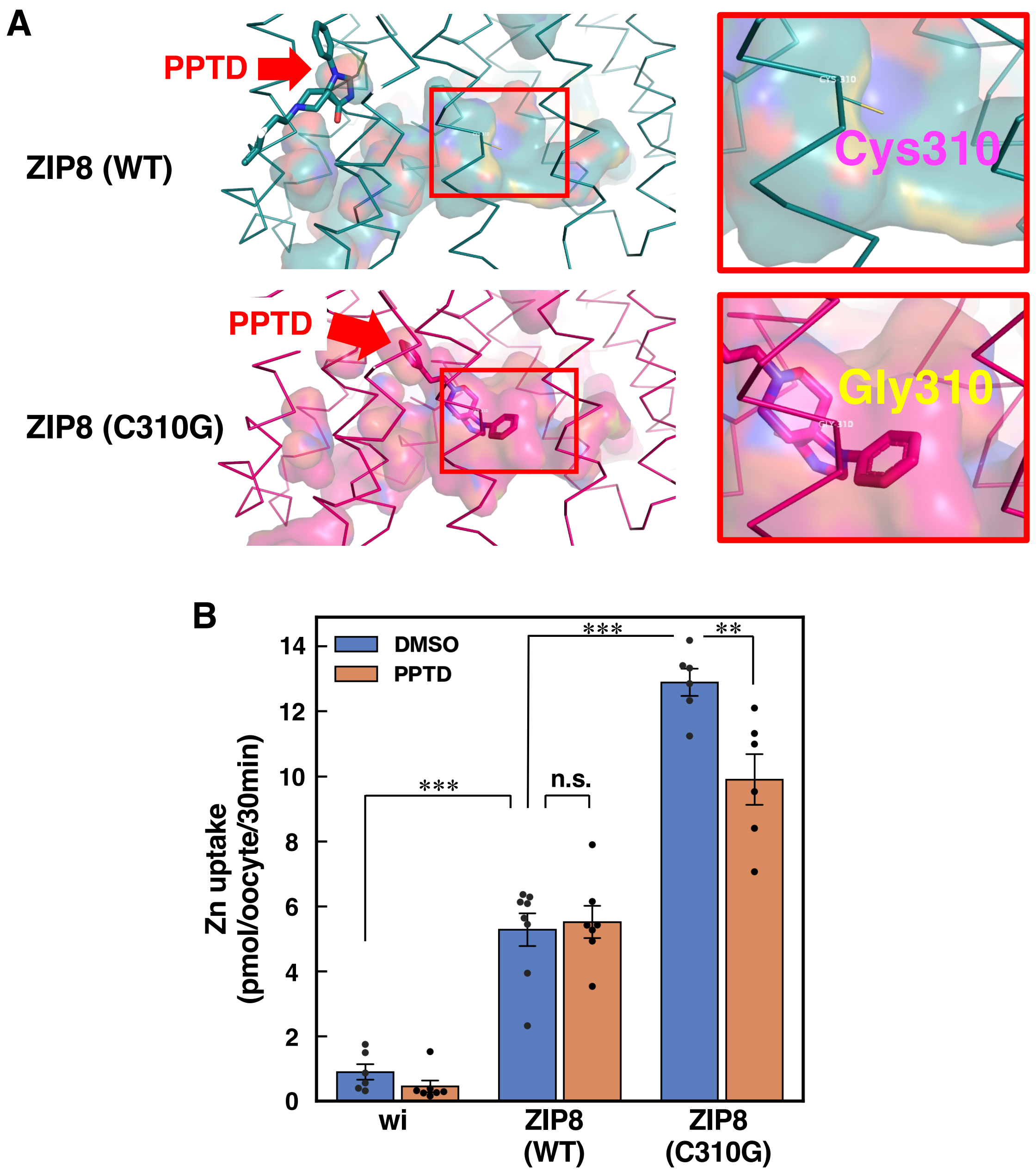


**Supplementary Fig. 6.**

**The pocket-creating C310G mutation in ZIP8 enhances zinc transport.**

**A** Computational modeling of human ZIP8 protein in complex with PPTD, comparing the wild-type (upper) and C310G mutant (lower) forms. Red arrows indicate PPTD binding sites. Enlarged views of the highlighted regions are shown in the right panels.

**B** *Xenopus* oocytes were injected with cRNA encoding either wild-type or C310G mutant human ZIP8, or with water (wi; control), and preincubated with 50 µM PPTD for 30 min, then incubated with 10 µM $ZnCl_2$ in the continued presence of PPTD for 30 min at 22 °C. Radioactivity was measured as described in the *Methods* section. Statistical analysis was performed using one-way ANOVA followed by Tukey's multiple comparison test. (**$p < 0.01$, ***$p < 0.001$). All data are shown as mean ± S.E.M. (n= 6 to 9 oocytes).

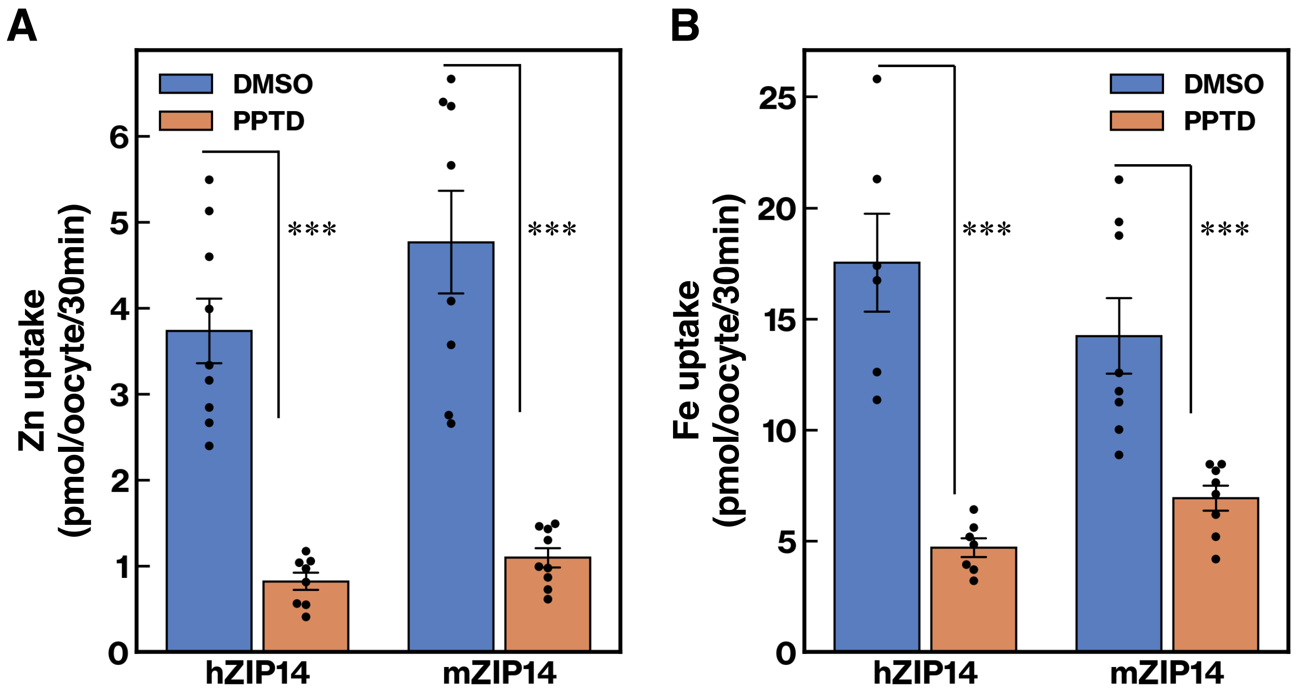


**Supplementary Fig. 7.**

**PPTD inhibits mouse ZIP14-mediated metal transport.**

*Xenopus* oocytes were injected with cRNA encoding either human ZIP14 (hZIP14), mouse ZIP14 (mZIP14), or with water (control). Oocytes were preincubated with 10 µM PPTD for 30 min, followed by incubation with 10 µM $Zn^{2+}$ (**A**) or $Fe^{2+}$ (**B**) in the continued presence of PPTD for an additional 30 min at 22 °C. Radioactivity was measured as described in the *Methods* section. The ZIP14-mediated transport activity in ZIP14-expressing oocytes was calculated by subtracting the transport activity in control oocytes. Statistical analysis was performed using one-way ANOVA followed by Tukey's multiple comparison test. (***$p < 0.001$). All data are shown as mean ± S.E.M. (n= 6 to 9 oocytes).

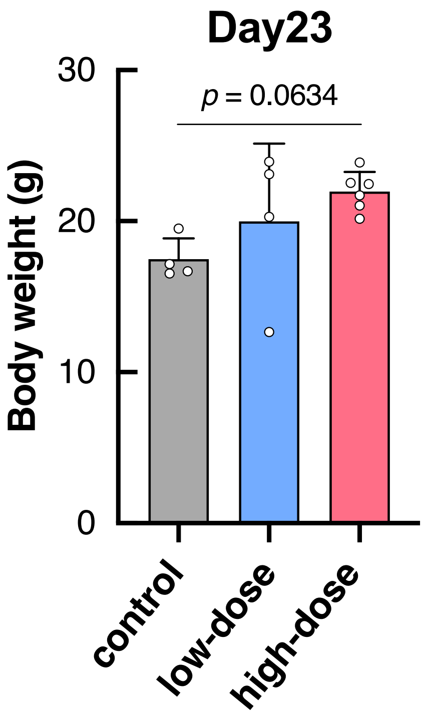


**Supplementary Fig. 8**

**PPTD administration tended to attenuate body weight loss.**

Effect of PPTD administration on body weight at day 23 in a cancer cachexia model. Mice were given PPTD in drinking water at concentrations of either 0.1 g/L (low-dose) or 1.0 g/L (high-dose).

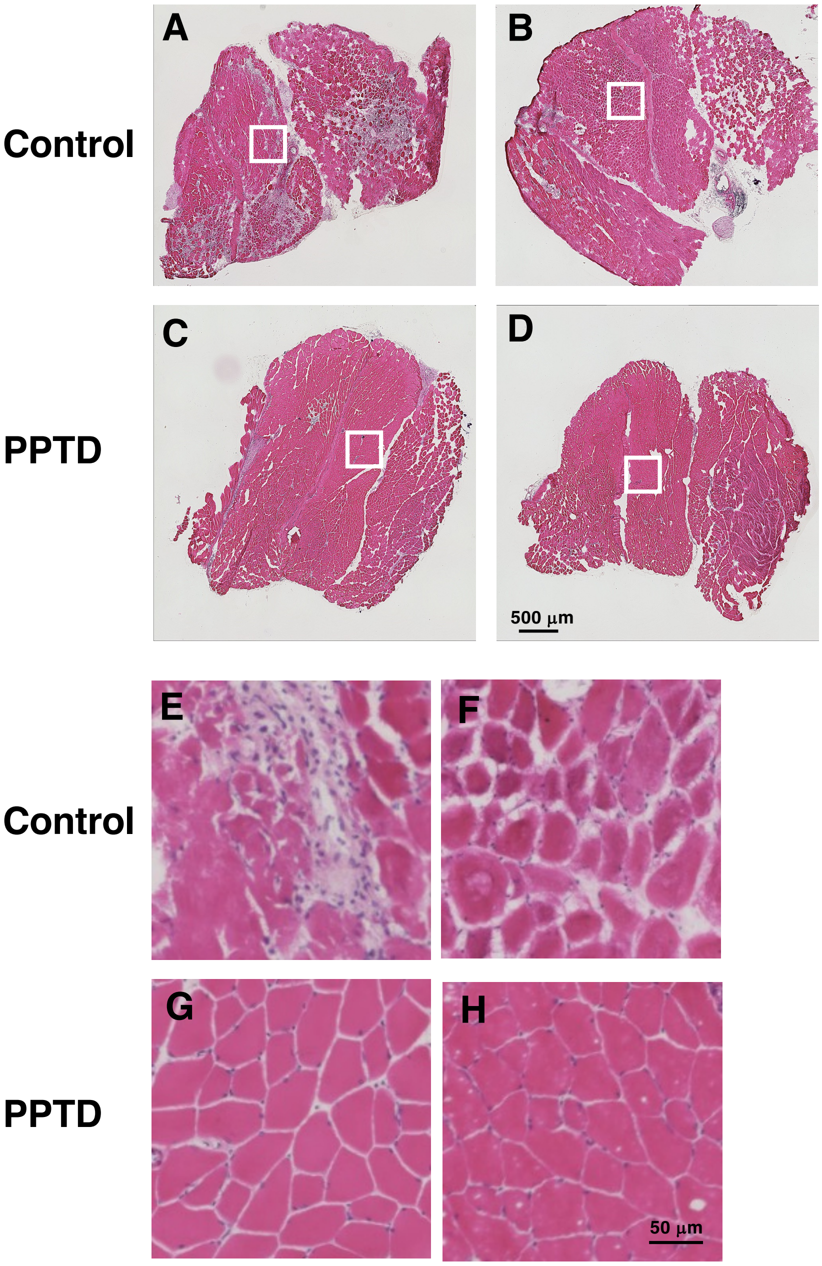


**Supplementary Fig. 9.**

**Effect of PPTD administration on skeletal muscle in a cancer cachexia mouse model.**

PPTD was administered ad libitum via drinking water to mice in a cancer cachexia model. Skeletal muscles of the lower limbs were collected 17 days after the start of treatment and analyzed by hematoxylin and eosin (HE) staining. Representative images from the control group (A, B) and the PPTD-treated group (C, D) are shown (2 mice per group). Panels E–H display magnified views of the white boxed areas in panels A–D, respectively. In the control group, loss of muscle fibers and infiltration of inflammatory cells were observed (A, E), along with variability in muscle fiber size, including both atrophic and hypertrophic fibers (B, F). In contrast, the PPTD-treated group exhibited densely packed muscle fibers with minimal inflammatory cell infiltration (C, D, G, H). These findings suggest that PPTD treatment suppresses inflammation in muscle tissue and mitigates muscle damage associated with cancer cachexia.

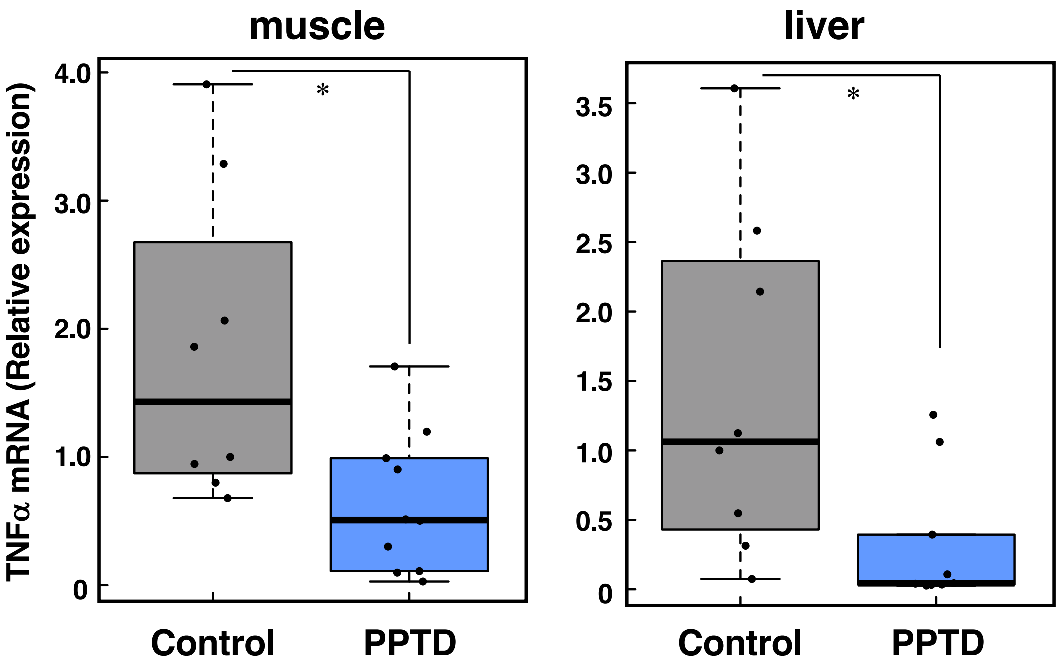


**Supplementary Fig. 10.**

**Effect of PPTD administration on *TNF-α* gene expression in a murine model of cancer cachexia.**

Mice were administered PPTD ad libitum via drinking water in a cancer cachexia model. Skeletal muscles of the lower limbs (left) and liver tissues (right) were collected before 17 days after the start of treatment, and *TNF-α* gene expression was analyzed by qPCR using specific primers (Supplementary Table 6) as described [1]. Box plots represent relative *TNF-a* expression levels with individual data points shown ($n$ = 8 to 10 per group). PPTD treatment suppressed *TNF-α* expression in both muscle and liver tissues in cancer cachexia mice. Statistical significance was evaluated using Welch's *t*-test ($*p < 0.05$).

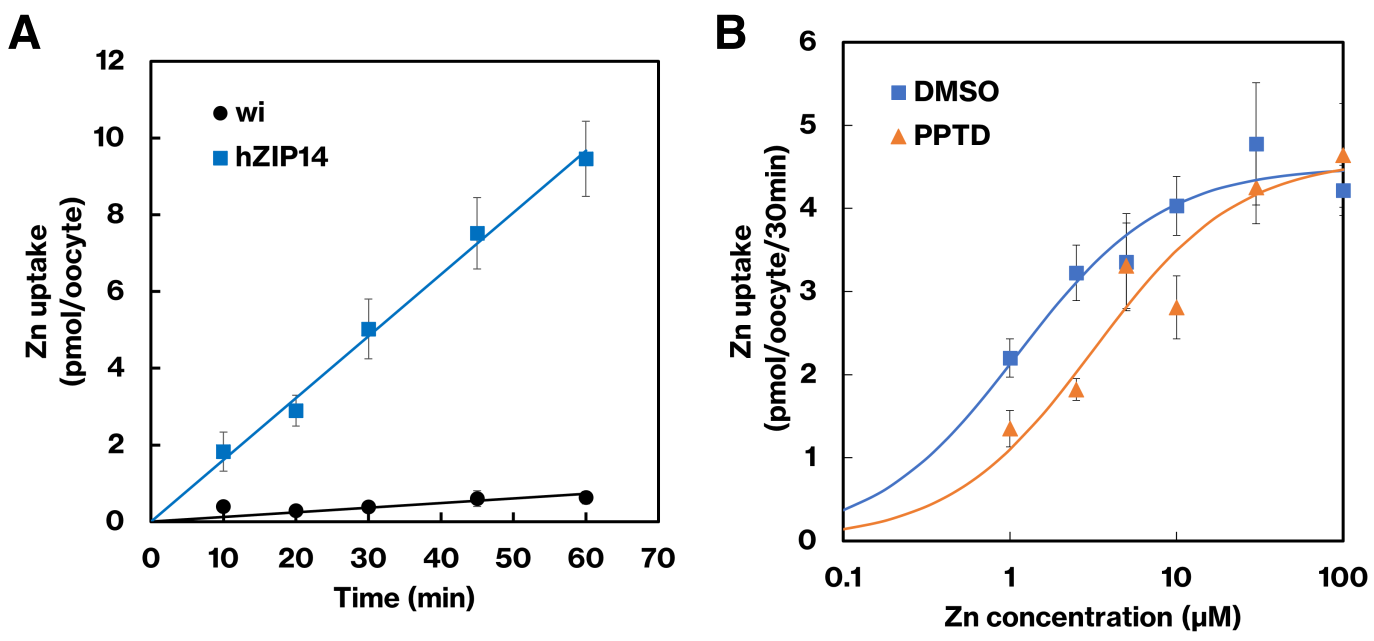


**Supplementary Fig. 11.**

**PPTD competitively inhibits ZIP14-mediated zinc transport.**

**A** Time course of zinc uptake by ZIP14. *Xenopus* oocytes injected with ZIP14 cRNA or water (wi: control) were incubated with 10 μM $ZnSO_4$ including a trace amount of radioactive tracer for the indicated times at 22 °C.

**B** PPTD competitively inhibits ZIP14-mediated zinc transport. Xenopus oocytes injected with ZIP14 cRNA or water (control) were incubated with indicated concentration of $ZnSO_4$ including a trace amount of radioactive tracer in the presence of 10 μM PPTD for 30 min at 22 °C. The ZIP14-mediated transport activity in ZIP14-expressing oocytes was calculated by subtracting the transport activity in control oocytes. All data are shown as mean ± S.E.M. (n= 6 to 9 oocytes).

#### Supplementary Tables

#### Supplementary Table 1. Comparison of $IC_{50}$ Values for Zinc and Iron Transport

| Transporting substrate | $IC_{50}$ | Two-sample Welch's t-test |
|---|---|---|
| Iron | 6.52μM ± 1.87 | *p* value = 0.85 |
| Zinc | 6.06μM ± 1.44 | |

$IC_{50}$ values for PPTD-mediated iron and zinc transport. Metal uptake assays were performed with 10 μM zinc or iron and 6 concentrations (0.03 to 50 μM) of PPTD. To estimate the $IC_{50}$ values, uptake values of 6-8 oocytes at each concentration were fitted to a 4-parameter logistic model using R package dr4pl (version 2.0.0). The results were expressed as mean ± S.E.M. (n=3 for zinc and n=5 for iron). Statistical significance was assessed by *Welch's t-test*, as described in the *Methods* section.

### Supplementary Table 2. Activity of PPTD-derivatives (N1-series)

| | Substitutive group | | |
|---|---|---|---|
| **Compound** | **R1** | **Relative Fe uptake (%)**<br>**(Fe uptake in the DMSO control was calculated as 100)** | **P-value**<br>**(vs. DMSO control)** |
| **stereoparent structure**<br>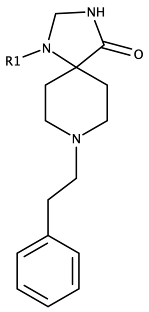 | | | |
| **PPTD** | 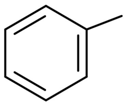 | **9.58±5.03** | **+ + + +** |
| **N1-2** | 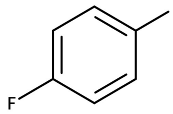 | **22.23±6.34** | **+ + + +** |
| **N1-1** | 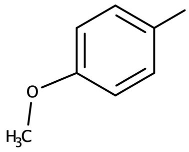 | **68.96±20.42** | **+** |
| **N1-3** | 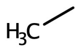 | **106.5±27.64** | **—** |

#### Supplementary Table 3. Activity of PPTD-derivatives (N3-series)

| | Substitutive group | | |
|---|---|---|---|
| **Compound** | **R3** | **Relative Fe uptake (%)** (Fe uptake in the DMSO control was calculated as 100) | **P-value** (vs. DMSO control) |
| stereoparent structure 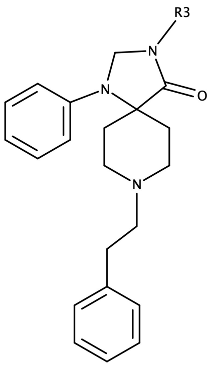 | | | |
| **PPTD** | **H** | **9.58±5.03** | **+ + + +** |
| **N3-1** | 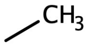 | **15.44±7.15** | **+ + + +** |
| **N3-2** | 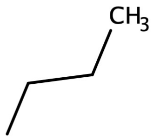 | **93.3±19.52** | **—** |
| **N3-3** | 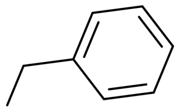 | **94.06±20.51** | **—** |

### Supplementary Table 4. Activity of PPTD-derivatives (N8-series)

| Compound | R2 | Relative Fe uptake (%)<br>(Fe uptake in the DMSO control was calculated as 100) | P-value<br>(vs. DMSO control) |
|---|---|---|---|
| stereoparent structure<br>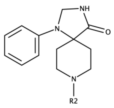 | | | |
| PPTD | 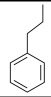 | 9.58±5.03 | + + + + |
| N8-3 | 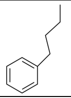 | 39.64±12.3 | + + + + |
| N8-2 | 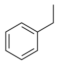 | 87.35±20.45 | — |
| N8-1 | H | 93.99±23.81 | — |
| N8-12 | 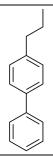 | 89.95±23.04 | — |
| N8-13 | 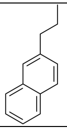 | 66.74±13.17 | — |
| N8-17 | 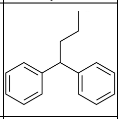 | 78.44±38.4 | — |
| N8-4 | 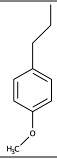 | 23.71±25.78 | + + + + |
| N8-9 | 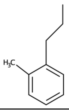 | 7.88±3.65 | + + + + |
| N8-6 | 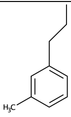 | 17.37±6.76 | + + + + |
| N8-8 | 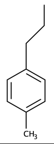 | 33.02±6.43 | + + + + |
| N8-10 | 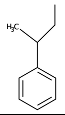 | 42.85±14.69 | + + + + |
| N8-18 | 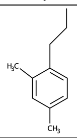 | 7.54±3.79 | + + + + |
| N8-19 | 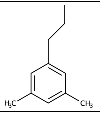 | 20.58±8.73 | + + + + |
| N8-20 | 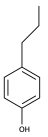 | 4.11±2.01 | + + + + |
| N8-21 | 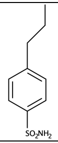 | 30.55±9.29 | + + + + |
| N8-22 | 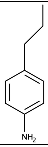 | 26.11±7.76 | + + + |
| N8-11 | 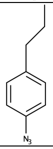 | 31.3±10.16 | + + + + |
| N8-23 | 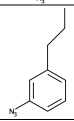 | 12.36±5.87 | + + + + |
| N8-24 | 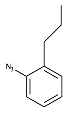 | 14.82±4.32 | + + + + |
| N8-5 | 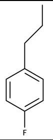 | 13.76±10.48 | + + + + |
| N8-14 | 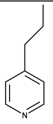 | 23.81±6.01 | + + + + |
| N8-15 | 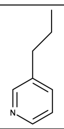 | 16.87±6.11 | + + + + |
| N8-16 | 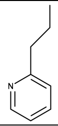 | 26.56±8.58 | + + + |
| N8-25 | 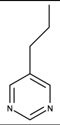 | 22.83±13.74 | + + + + |
| N8-7 | 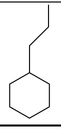 | 17.1±10.87 | + + + + |

#### Supplementary Table 5. Computational Prediction of the High-Affinity Interaction Between PPTD and Human ZIP14 Based on PLIP and AlphaFold3 Analyses

| Interaction | Residues |
|---|---|
| Hydrophobic Interactions | Phe 234 |
| | Glu 238 |
| | Leu 346 |
| | His 347 |
| Hydrogen Bonds | Ser 343 |
| Salt Bridges | Asp 384 |
| | Asp 443 |
| pi-Stacking | His 380 |

The corresponding structural model of the PPTD–ZIP14 complex, generated based on PLIP analysis in conjunction with AlphaFold3 predictions, is presented in Fig. 5C.

#### Supplementary Table 6. Primer Sequence used for qPCR Analyses

| Gene | Forward (5'→3') | Reverse (5'→3') |
|---|---|---|
| mouse TNF$\alpha$ | CAGGCGGTGCCTATGTCTC | CGATCACCCCGAAGTTCAGTAG |
| mouse 18s rRNA | AGTCCCTGCCCTTTGTACACA | GATCCGAGGGCCTCACTAAAC |

#### Legends for Supplementary Movies

**Supplementary Movie 1.**
**Three-dimensional animation of monomer human ZIP14 and PPTD complexes.**
This 3D animation shows the predicted structure of monomer human ZIP14 bound to PPTD, as modeled using AlphaFold3 (corresponding to Fig. 5A left). magenta: PPTD.

**Supplementary Movie 2.**
**Three-dimensional animation of dimerized human ZIP14 and PPTD complexes.**
This 3D animation shows the predicted structure of dimerized human ZIP14 bound to PPTD, as modeled using AlphaFold3 (corresponding to Fig. 5A right). magenta: PPTD.

**Supplementary Movie 3.**
**Three-dimensional overlay of dimerized human ZIP14–PPTD and ZIP14–zinc complexes.**
This 3D animation shows the predicted structural overlay of dimerized human ZIP14 bound to PPTD and to zinc, as modeled using AlphaFold3 (corresponding to Fig. 5B). The comparison highlights the overlapping ligand-binding sites within the human ZIP14 dimer. orange: PPTD, blue: amino residues, purple sphere: zinc.

**Supplementary Movie 4.**
**Three-dimensional animation of the interaction between human ZIP14 and PPTD.**
This 3D animation depicts the predicted interaction between human ZIP14 and PPTD, as modeled using the Protein–Ligand Interaction Profiler and AlphaFold3 (corresponding to Fig. 5C and Supplementary Table 5). orange: PPTD, blue: amino residues, green dot line: pi-Stacking, yellow dot line: salt bridges, gray dot line: hydrophobic interactions, blue line: hydrogen bond.

**Supplementary Movie 5.**
**Three-dimensional animation of the PPTD-binding pocket within human ZIP14.**
This 3D animation depicts the PPTD-binding pocket within human ZIP14 (corresponding to Fig. 6A upper). green: PPTD

**Supplementary Movie 6.**
**Three-dimensional animation of the PPTD-binding pocket in human ZIP14 with the substitution of Serine 343 with Tyrosine (S343Y).**
This 3D animation depicts the PPTD-binding pocket in human ZIP14 with the substitution of Serine 343 with Tyrosine (S343Y) reducing the size of the PPTD-binding pocket within ZIP14 (corresponding to Fig. 6A lower). yellow: PPTD

**Supplementary Movie 7.**
**PPTD administration improves locomotor activity in a murine cachexia model.**
Representative videos of cachectic mice on day 16, with (top) or without (bottom) PPTD treatment, are shown at three times the normal speed.
